# Integrating genomics and multi-platform metabolomics enables metabolite QTL detection in breeding-relevant apple germplasm

**DOI:** 10.1101/2021.02.18.431481

**Authors:** Emma A. Bilbrey, Kathryn Williamson, Emmanuel Hatzakis, Diane Doud Miller, Jonathan Fresnedo-Ramírez, Jessica L. Cooperstone

**Affiliations:** Department of Horticulture and Crop Science, The Ohio State University, Columbus, OH, USA; Department of Food Science and Technology, The Ohio State University, Columbus, OH, USA; Department of Horticulture and Crop Science, The Ohio State University, Wooster, OH, USA

**Keywords:** apple breeding, chlorogenic acid, genome-wide association study (GWAS), *Malus* x *domestica* (apple), omics, pedigree-based analysis (PBA), quantitative trait loci (QTL)

## Abstract

**Research Conducted:** Apple (Malus × domestica) has commercial and nutritional value, but breeding constraints of tree crops limit varietal improvement. Marker-assisted selection minimizes these drawbacks, but breeders lack applications for targeting fruit phytochemicals. To understand genotype-phytochemical associations in apples, we have developed a high-throughput integration strategy for genomic and multi-platform metabolomics data.

**Methods:** 124 apple genotypes, including members of three pedigree-connected breeding families alongside diverse cultivars and wild selections, were genotyped and phenotyped. Metabolite genome-wide association studies (mGWAS) were conducted with 10,000 single nucleotide polymorphisms and phenotypic data acquired via LC-MS and ^1^H NMR untargeted metabolomics. Putative metabolite quantitative trait loci (mQTL) were then validated via pedigree-based analyses (PBA).

**Key Results:** Using our developed method, 519, 726, and 177 putative mQTL were detected in LC-MS positive and negative ionization modes and NMR, respectively. mQTL were indicated on each chromosome, with hotspots on linkage groups 16 and 17. A chlorogenic acid mQTL was discovered on chromosome 17 via mGWAS and validated with a two-step PBA, enabling discovery of novel candidate gene-metabolite relationships.

**Main Conclusion:** Complementary data from three metabolomics approaches and dual genomics analyses increased confidence in validity of compound annotation and mQTL detection. Our platform demonstrates the utility of multi-omics integration to advance data-driven, phytochemicalbased plant breeding.

## Introduction

Apples *(Malus × domestica* Borkh.) are eaten throughout the world and the most consumed fruit in the US (USDA Economic Research Service, 2017). Barriers to apple breeding, including long juvenile period, self-incompatibility, and heterozygous genomes, necessitate a breeding cycle based on marker-assisted selection to make efficient progress by allowing breeders to choose parents with high heritability for desirable traits and implement early-seedling selection. Traits important in apple breeding, such as nutrition, flavor, texture, disease resistance, and post-harvest quality, are collective phenotypes largely impacted by the phytochemical makeup of apple fruits (Boyer | & Liu, 2004; Pérez |& Sanz, 2008; Hyson, 2011; Sun et al., 2017; Vondráková et al., 2020). However, metabolite-based breeding strategies for apple improvement are lacking.

To determine genetic regions associated with phenotypic traits in apple, including metabolite abundance, quantitative trait loci (QTL) mapping has traditionally been performed with bi-parental families (Dunemann et al., 2009; Chagné et al., 2012; Khan et al., 2012; Verdu et al., 2014; Gutierrez et al., 2018; Christeller et al., 2019). However, due to the heavily heterozygous nature of the apple genome and diversity available in wild and cultivated species, bi-parental populations hamper characterization of possible alleles. Selection based on information about two to four alleles present in a bi-parental population then leads to loss of efficiency and genetic erosion when other genotypes are ignored (van de Weg et al., 2004). Loci that are characterized in one mapping population are commonly not transferrable to other families. Furthermore, analysis using segregating bi-parental populations is often impractical due to the long juvenile period that would stretch the production of segregating populations to many decades. As a result, it is necessary to move beyond bi-parental populations in apple trait loci analysis (Peace et al., 2019).

Generating a metabolite-based apple breeding strategy requires us to develop our foundational understanding of (1) the phytochemical profiles that exist in apple germplasm and (2) the genetics underlying this variability. Global phytochemical assessment via untargeted metabolomics approaches can be obtained by using high-resolution mass spectrometry (MS) as well as nuclear magnetic resonance (NMR) spectroscopy analyses. Genetic data can be gathered with single nucleotide polymorphism (SNP) arrays to characterize genotypes of chosen apple varieties. The larger challenge comes with (3) the integration of the genomic and metabolomic datasets to determine the genotype-metabolite relationships that are the basis of marker development. Moving from analyzing genetic control of a few traits to the thousands of phenotypes contained in a metabolomics dataset, requires a high-throughput pipeline tailored to perennial crops, such as apple.

To this end, we have developed a scheme for integrating SNP array data with multi-platform, untargeted metabolomics datasets to identify metabolite QTL (mQTL) in breeding-relevant apple germplasm (Fig. 1). The method uses metabolite genome-wide association studies (mGWAS) to detect putative mQTL. To reduce false positives and identify mQTL with robust signal across available apple germplasm, three mGWAS iterations are conducted for three nested population sets, each adding more complexity: (1) three sets of progenies, (2) progenies plus pedigree-related individuals, and (3) progenies, pedigree-related individuals, and additional heritage and wild selections. mQTL have to be detected in each of these three groupings to protect against spurious correlations. Three separate, complementary metabolomics platforms, MS in positive and negative ionization modes and NMR, are used to aid in mQTL detection and facilitate metabolomic feature identification. Metabolites of interest are then passed to a more rigorous pedigree-based analysis (PBA) for mQTL validation through linkage analysis and identity-by-descent (IBD). Findings are then contextualized according to the current annotated apple genome.

**Fig. 1.**
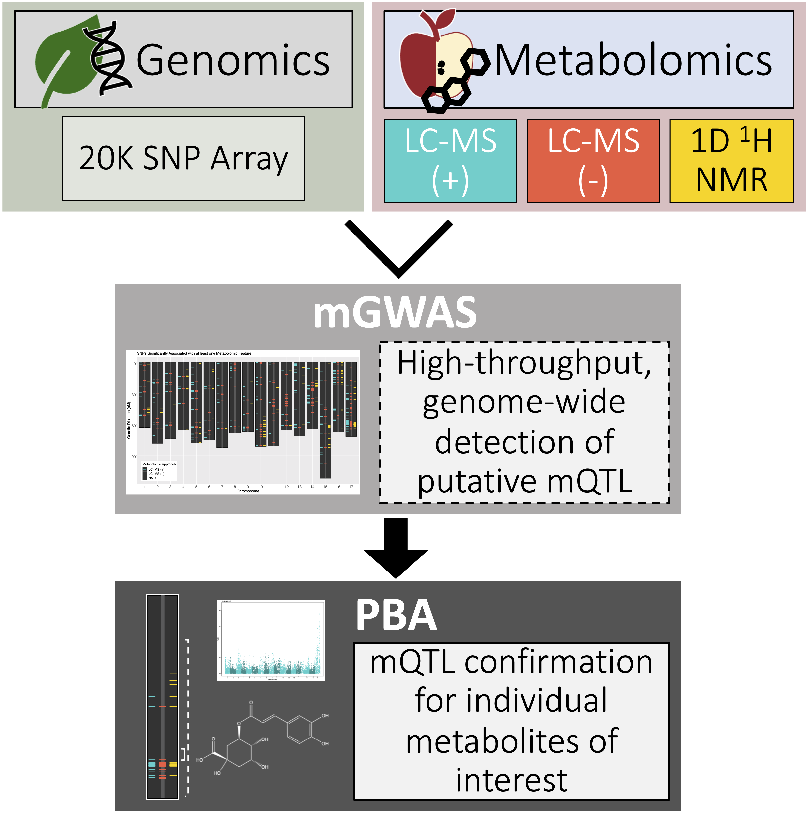
Our platform for detecting metabolite quantitative trait loci (mQTL) is based on integrating SNP array data and multi-platform metabolomics via metabolite genome-wide association studies (mGWAS) then confirming mQTL for metabolites of interest with pedigree-based analyses (PBA).

## Materials and Methods

### Apple Germplasm

Apple genotypes (n=124) were chosen for the study based on the expected diversity of their genetic and metabolic profiles, as well as commercial interest. Three sets of experimental progenies from a grower participatory breeding program, including ‘Honeycrisp’ × ‘Fuji’ (HC×FJ, n=28), ‘Goldrush’ × ‘Sweet 16’ (GR×S16, n=28), and ‘Honeycrisp’ × ‘MSH 10-1’ (HC×M10, n=19), with varied flavor and texture profiles were selected along with 23 members of their pedigree-connected families (Sup. Fig. S1). These three families (n=98) include commercial and heritage apple varieties, *M. floribunda* Siebold ex Van Houtte, and advanced selections. Wild accessions (n=11) from Central Asian M. *niedzwetzkyana* (Dieck) Langenf and M. *sieversii* (Lebed) Roem along with additional apples (n=15) with traits of breeding interest were also included to capture the wide variety of apple germplasm. All apples and metadata including collection location, date, and harvest-to-storage protocol can be found in Sup. Table S1 and Methods S1.

### Genotypic Information

DNA extraction from leaf tissue was carried out using the Omega E-Z 96 Plant DNA Kit (Omega Bio-tek, Inc. Norcross, GA, USA). Samples were then genotyped at Michigan State University using an apple 20K Infinium^®^ SNP array (Bianco et al., 2014). SNP calling and filtering was performed using GenomeStudio version 2.0.4 (Illumina Inc., San Diego, CA, USA; http://www.illumina.com). Marker order was determined by integrating markers with the apple integrated genetic linkage map (iGLMap) (Di Pierro et al., 2016).

Pedigree relationships were verified using 1,648 SNPs in FRANz 2.0 (Riester et al., 2009), as in Fresnedo-Ramírez et al. (2015). SNPs were filtered based on a minor allele frequency (MAF) minimum of 0.10 and missingness of less than 1%. Parentage was corrected for accessions with high posterior probability (>0.95) of distinct parentage.

Less stringent filtering was applied to the marker dataset to determine SNPs to use for mGWAS. Markers were first matched to those included in the iGLMap, giving 15,260 SNPs. These were then filtered for minimum MAF > 0.05 and missingness of 5%. This resulted in 10,294 polymorphic, genome-wide markers to be used for mGWAS analyses (Sup. Table S2).

A third set of parameters were used to filter markers to be used in PBA. Markers were kept with MAF > 0.07 and missingness less than 10% in the pedigree members in order to yield the maximum number of informative markers that also matched with those represented in the iGLMap. The markers were analyzed in FlexQTL™ v.099130 at the Ohio Supercomputer Center (Ohio Supercomputer Center, 1987) to check for double recombinations and an excessive number of genotyping inconsistencies. Markers were removed with >3 genotyping inconsistencies as well as any obvious double recombinants, resulting in a final set of 6,034 markers for use in the PBA routines.

### Metabolomic Phenotyping

Methanolic extracts were prepared for each apple selection and subsequently analyzed via ultra-high performance liquid chromatography-quadrupole time-of-flight mass spectrometry (UHPLC-QTOF-MS) in both electrospray ionization (ESI) positive (+) and negative (-) modes as well as 1D ^1^H NMR experiments on a 700.13 MHz instrument. LC-MS raw spectral data was deconvoluted using MZmine2.51 (Pluskal et al., 2010). Pooled quality control (QC) samples were used to confirm data quality. Process blank analyses allowed detection and removal of any residues or contamination with compounds from materials used in the extractions. NMR spectra were baseline and phase corrected and adjusted to trimethylsilylpro pionic acid (δ=0) using the Topspin software package provided by Brüker Biospin. NMR spectral post-processing was performed using R package mrbin v1.2.9001 (Klein, 2020). Complete details on extraction, sample analysis, and data processing are available in Sup. Methods S2-S4 and Tables S3-S5.

Remaining features from LC-MS and NMR analyses were log2-transformed then analyzed in R 3.6.2 (R Development Core Team, 2008). Unsupervised principal components analyses (PCA) were conducted for each of the three final metabolomics datasets: LC-MS (+), LC-MS (-), and NMR.

### Feature identification

Data-dependent UHPLC-QTOF-MS/MS experiments were performed on pooled QC samples in both ionization modes to obtain fragmentation data for putative identity generation via library searches within the Global Natural Products (GNPS) platform (Wang et al., 2016) as well as the Human Metabolome Database (HMDB) (Wishart et al., 2018). Additional information can be found in Sup. Methods S5-S6.

A chlorogenic acid authentic standard was purchased for targeted MS/MS analysis. Resultant spectra from the standard were compared with the accurate mass, retention time, and fragmentation pattern of the peak of interest in a pooled QC sample. The standard was also used in spike experiments analyzed via 1D and 2D NMR to confirm peak identities in the NMR spectra.

## Omics Integration for mQTL Detection

### Metabolite genome-wide association studies (mGWAS)

To integrate genomics and metabolomics data for mQTL detection, mGWAS analyses were conducted in which each metabolomic feature (n=10,325) was considered a phenotype and was examined for association with each SNP (n=10,294). Because of the large number of phenotypes, individual model optimization was not possible, so generalized strategies for analyses were developed. Within each genomics-metabolomics platform combination, three separate analyses of certain apple genotypes were conducted: progenies only (Progeny, n=75), progenies plus pedigree-related individuals (Pedigree, n=98), and a full, diverse analysis of all individuals for which genomics and metabolomics data were collected (Diverse, n=124).

We developed a workflow to assess SNP-feature associations within three population sets (Fig. 2b) for each of the three metabolomics datasets (LC-MS (+), LC-MS (-), and NMR) (Fig. 2). This resulted in a total of nine multivariate sets of mGWAS analyses (Fig. 2a) performed using the R package rrBLUP v4.6.1 (Endelman, 2011). In order to model known pedigree and realized genetic relatedness, R package AGH-matrix v1.0.2 (Amadeu et al., 2016) was used to generate an H matrix (Legarra et al., 2009), which combines theoretical additive relatedness estimations based on pedigree connections with realized genomic relatedness estimations from molecular markers (Sup. Tables S11-S12). To correct for additional structure within the populations analyzed, PCA scree plots of SNP data were examined to determine the appropriate number of principal components to include in the analyses (Progeny: 3, Pedigree: 6, Diverse: 10) (Sup. Fig. S2). Unix batch scripts containing R code were executed at the Ohio Supercomputer Center (Ohio Supercomputer Center, 1987). From each of the nine mGWAS versions, results for separate chromosomes were collated to produce a single data frame of -log_10_(P) values for each pairwise SNP-feature association, resulting in nine unique SNP-by-feature data frames – three per metabolomics platform.

**Fig. 2.**
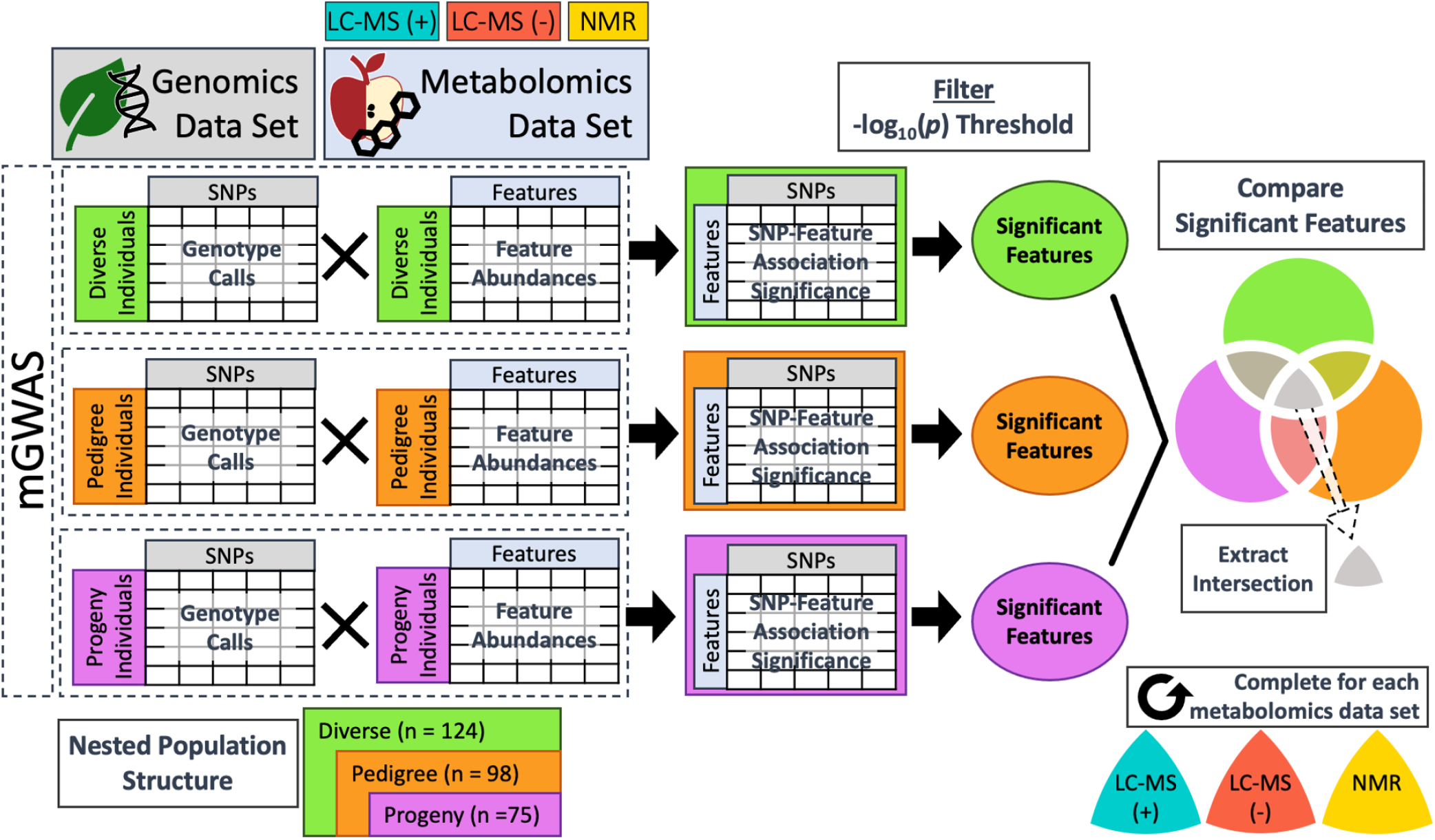
Workflow for integration of one metabolomics dataset with genomics data. This workflow was applied to each of the three metabolomics datasets. Three separate mGWAS analyses (**a**) with different subsets of individuals (**b**) were conducted in order to detect real SNP-feature associations present across diverse germplasm and in segregating progeny. This was achieved by filtering results of each for strong signal (**c**) and then comparing the results from the three populations (**d**). Features represented in the intersection (**e**) were extracted for further analysis and identification. This resulted in a corresponding collection of overlapping significant features (**f**) for each of the three metabolomic datasets.

SNP-feature associations were filtered for significance with – log_10_(P) threshold (Fig. 2c). For LC-MS (+) and (-) datasets, a threshold of ≥ 4 (P<.0001) was used for the progeny results and ≥ 5 (P<.00001) for the pedigree and diverse results. A ≥ 4 filter was used for each of three populations of NMR analyses due to the fewer comparisons relative to the LC-MS datasets. Values were not subjected to a multiple test correction, but significance thresholds displayed on all Manhattan plots correspond to P = .05 with a false discovery rate (FDR) correction for that feature.

From the significance-filtered mGWAS results, sets of features were compared via Venn diagrams (Progeny ⋂ Pedigree ⋂ Diverse) (Fig. 2d). The intersection (Progeny U Pedigree U Diverse) was then extracted (Fig. 2e). This process resulted in three core datasets, one per metabolomics platform, of -log_10_(P) values for features significantly associated with at least one SNP in each population set (Fig. 2f). The features in these core datasets were then considered as having putative mQTL.

To advance with these three sets of putative mQTL, additional approaches for broad data visualization were adopted, enabling the focus of further analyses. To investigate the distribution of these mQTL within each chromosome, a composite map of mQTL across the genome using all three metabolomic platforms was created. Additionally, to understand if many disparate SNPs were eliciting signal or if a few SNPs were associated with many metabolomic features, plots were constructed to visualize the number of significantly associated features per SNP within each chromosome.

As an illustrative example of the mGWAS workflow, a significant feature from each metabolomics platform was identified as chlorogenic acid (5-caffeoylquinic acid), and the phenotypic measurements were used as inputs in the PBA pipeline. This metabolite of the phenylpropanoid biosynthesis pathway had significant associations across the three metabolomics datasets and strong putative identification based on an MS/MS database match, which was confirmed by comparison with an authentic standard in both MS/MS and 1D and 2D NMR experiments.

### Pedigree-based Analysis (PBA)

Pedigree-based analyses (PBA) under a Bayesian framework were conducted using FlexQTL™ version 0.99130 for Linux (Bink 2002; Bink et al. 2002) (Sup. Methods S7). The PBA method incorporated pedigree relationships, genotype calls (Sup. Table S9), genetic linkage information in the form of genetic distances (recombination between markers) based on the iGLMap (Di Pierro et al., 2016), and log_2_-transformed metabolite abundance data. The 98 pedigree-related individuals were used in this analysis performed using a model that estimated additive and dominance effects within the pedigree. Each genomewide analysis was performed in triplicate, using at least 50,000 iterations in the Markov chain Monte Carlo (MCMC) procedure, with 1,000 burn-in iterations, sampling each 50^th^ chain to yield 1,000 effective samples for statistical analyses. The measurements for chlorogenic acid from each of the metabolomics datasets were analyzed for linkage separately. For NMR, chlorogenic acid signal is represented by several bins, so the bin with the highest -log_10_(P) value from the mGWAS, 2.15-2.14 ppm, was chosen for analysis. mQTL signals were considered positive when twice the natural log of the Bayes factor (2ln(BF)) was found to be 2-5, strong when 5-10, and decisive at >10. Narrow sense heritability was calculated by dividing the phenotypic variance by the sum of the weighted additive genetic variance and the sample residual variance. Subsequently, targeted IBD analysis was performed in FlexQTL™ to distinguish haplotypes and genotype (i.e., QQ, *Qq,* or qq) the mQTL region to estimate breeding values. Here, the mQTL was divided into bins of 1 cM, and the probability that a given locus contained a genetic component influencing chlorogenic acid abundance was measured via PBA in FlexQTL™.

In order to contextualize the presence of a chlorogenic acid mQTL, the characterized apple genome (GDDH13v1.1) was examined in the area surrounding the mQTL (Daccord et al., 2017; Jung et al., 2019). Genes encoding enzymes connected with chlorogenic acid production were noted.

## Results

### Pedigree Confirmed and Revised

Analysis of adherence to expected pedigree relationships indicated 18 individuals with disparate parentage. Due to phenotypic interest, six of these remained in the mGWAS under the Diverse category but were not considered part of the pedigree if both parents were unknown (Sup. Table S1). Additionally, the traditional lineage of the advanced selection ‘Co-op 17’ was revised to include ‘Crandall’ as a progenitor.

### Metabolomic Profiling Demonstrates Apple Metabolome Variety

Data processing of untargeted metabolomic analyses of the apple extracts resulted in 4,866 molecular features for LC-MS (+), 4,703 for LC-MS (-), and 756 bins for NMR (Sup. Tables S6-S8). No samples were found to be outliers when examining boxplots of each sample (Sup. Fig. S3). Good data quality was indicated for LC-MS experiments due to tight clustering of regularly injected pooled QCs when each ionization mode was examined separately via PCA (Sup. Fig. S4).

PCA showed metabolomic variation across genotypes in all metabolomic datasets (Sup. Fig. S5). The spread of points between the classifications of progeny, pedigree-connected, and other diverse selections confirmed that the apples selected for this study have metabolic variety.

### mGWAS Pipeline for Prioritization

The nine result matrices from the mGWAS analyses (Sup. Tables S13-S21) were filtered for significance (Fig. 3a). Then, extracting the intersection of the significant associations for the three population sets within each metabolomics dataset resulted in 519 LC-MS (+), 726 LC-MS (-), and 177 NMR features with putative mQTL (Fig. 3b-d). These features are listed in Sup. Table S22. Potential mQTL from the three metabolomics platforms were discovered on each of the 17 chromosomes (Fig. 4). An even distribution of mQTL across the 17 chromosomes would indicate 5.9% would be present per chromosome. We identified linkage group (LG) 16 as a hotspot because 54-60% of the mQTL detected per metabolomic approach (LC-MS (+) 284, LC-MS (-) 443, NMR 95) are located there. Linkage group 17 had an over-representation of mQTL as well: LC-MS (+) 97, LC-MS (-) 131, and NMR 7. LC-MS approaches provided more signals for mQTL on LG 17 compared to NMR – 18-19% of total mQTL versus 4% for NMR.

**Fig. 3.**
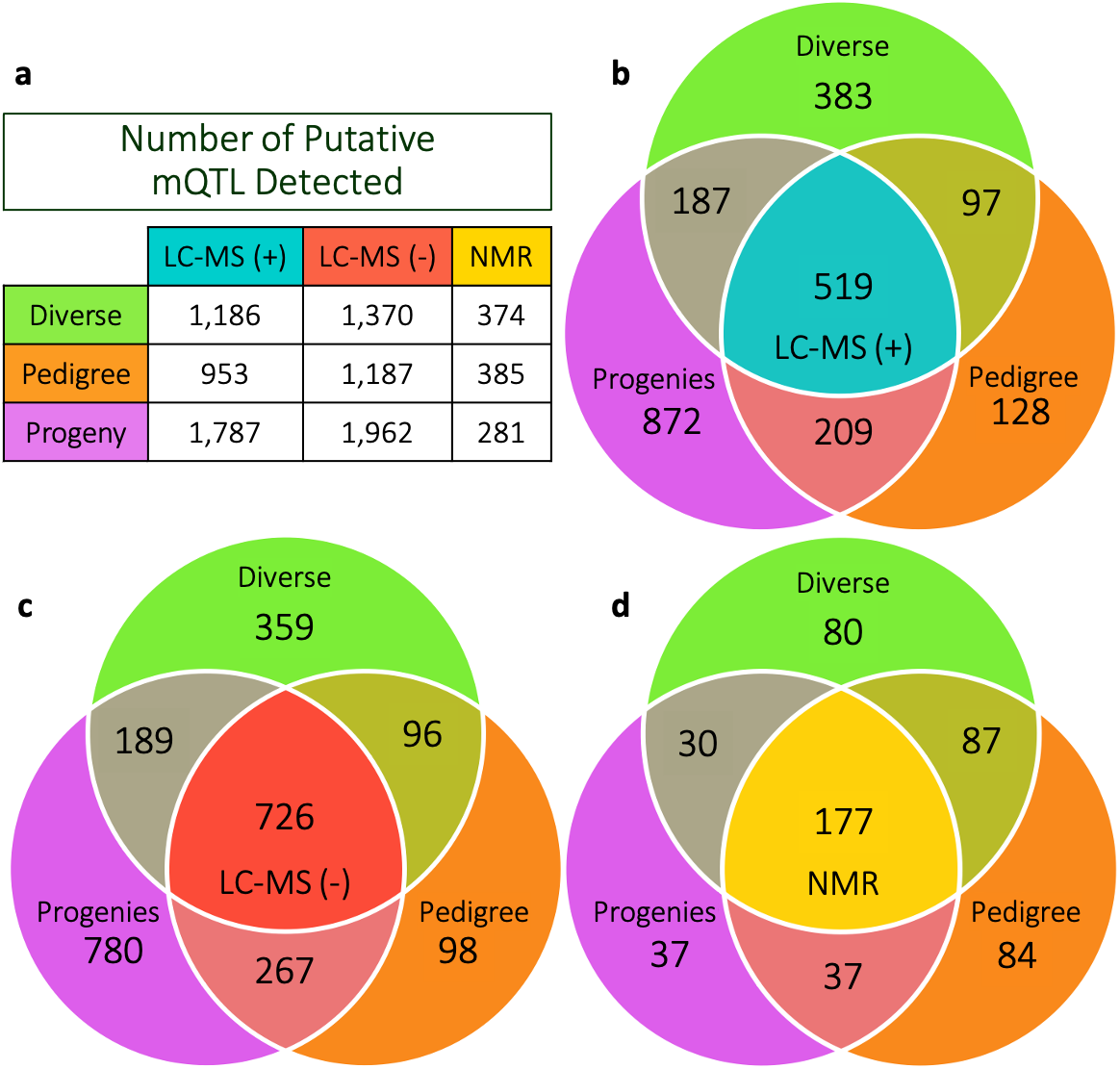
(**a**) The total counts of metabolomic features with SNP associations above significance thresholds for each population set in each metabolomics platform. The features were then compared within each metabolomics dataset to extract the intersection: a list of those that were significant in all three populations. Corresponding Venn diagrams for (**b**) LC-MS (+), (**c**) LC-MS (-), and (**d**) NMR metabolomic features are shown with the intersection containing the number of features that passed on for further analysis.

**Fig. 4.**
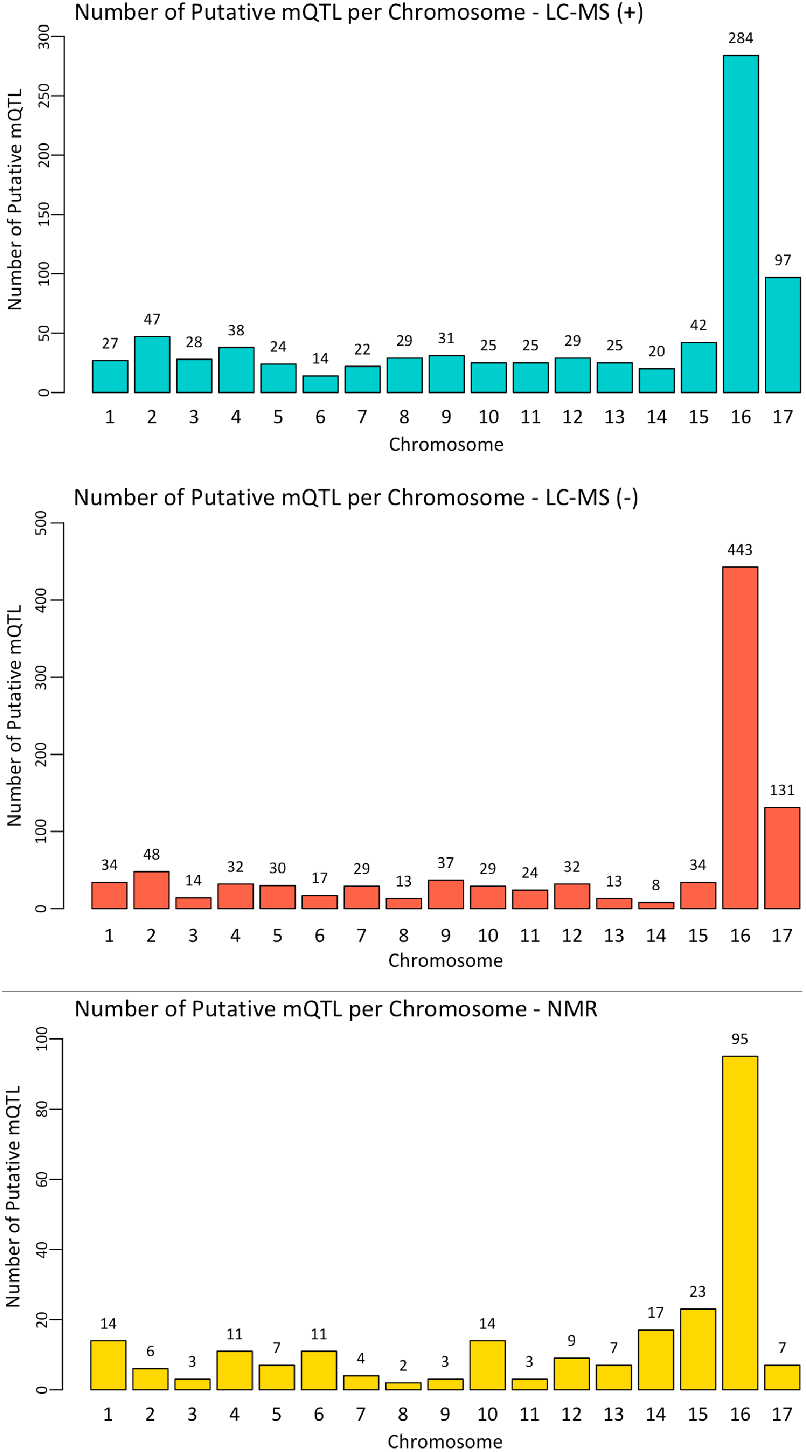
Bar plots showing the number of putative mQTL detected per chromosome via mGWAS for each metabolomics platform.

Simultaneous visual assessment of genomic areas that housed significantly associated SNPs using the three metabolomics datasets was achieved by creating a composite mQTL chromosome map (Fig. 5). Here, hotspots on the top of LG 16 and bottom of 17 are easily visible. Certain areas of the genome elicited mQTL signals for compounds in each of the metabolomic datasets (e.g., top of LG 16 and bottom of 17). Others were only significantly associated with one (e.g., middle of LG 13) or two of the metabolomics techniques (e.g., top of LGs 2 and 3). Supplementary Figs. S6 and S7 detail the number of features significantly associated with each SNP in LG 16 and 17, respectively. Clear regions of fruit phytochemical control were evident, as a set of SNPs on the top of LG 16 were significantly associated with hundreds of features across the three analysis methods. Particularly, three SNPs (13681, 13685, and 13675) maintained top positions for LC-MS (+), (-), and NMR (SNP name reference is available in Sup. S10). Similarly, on LG 17, a large number of signals clustered towards the bottom of the chromosome. Once again, parallel results were seen across each metabolomic approach with top ranked SNPs being 15109, 15123, and 15133.

**Fig. 5.**
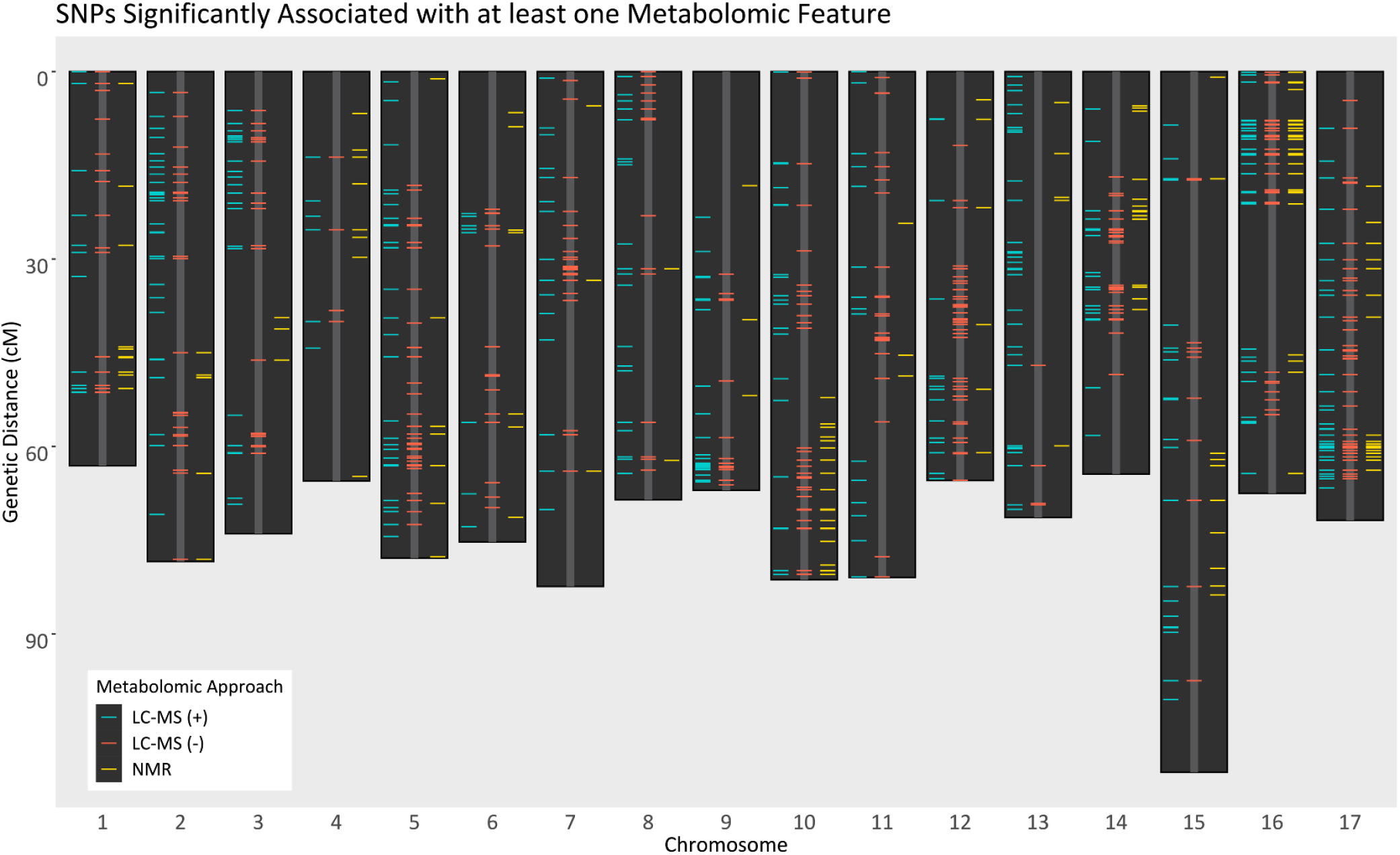
Composite mQTL chromosome map of the 17 apple chromosomes. Horizontal lines indicate the locations of each SNP found to have a significant association with at least one metabolomic feature in the filtered and intersected mGWAS results. Lines are colored based on the origin of the metabolomic feature.

### Chlorogenic Acid mQTL Proposed by mGWAS Pipeline and Validated with Two-Step PBA

The pipeline described was able to discover the relationship between chlorogenic acid and an mQTL on LG 17. Data-dependent MS/MS analyses were mined for features of final significance from the mGWAS analyses. This analysis yielded spectral matches to chlorogenic acid in GNPS (Wang et al., 2016) for both LC-MS (+) and (-). Identity was confirmed for peaks in both LC-MS and NMR datasets using authentic standards—a Level 1 annotation according to the Metabolomics Standard Initiative (Sumner et al., 2007).

Manhattan plots of mGWAS results for the chlorogenic acid features each showed significant signals (FDR-corrected P<.05) on LG 17 along with a suggestive signal on LG 3 (Fig. 6). SNP 15109 (Chr17:27,490,016) was indicated as the most significant locus in the LG 17 signal from the mGWAS (Fig. 7b).

**Fig. 6.**
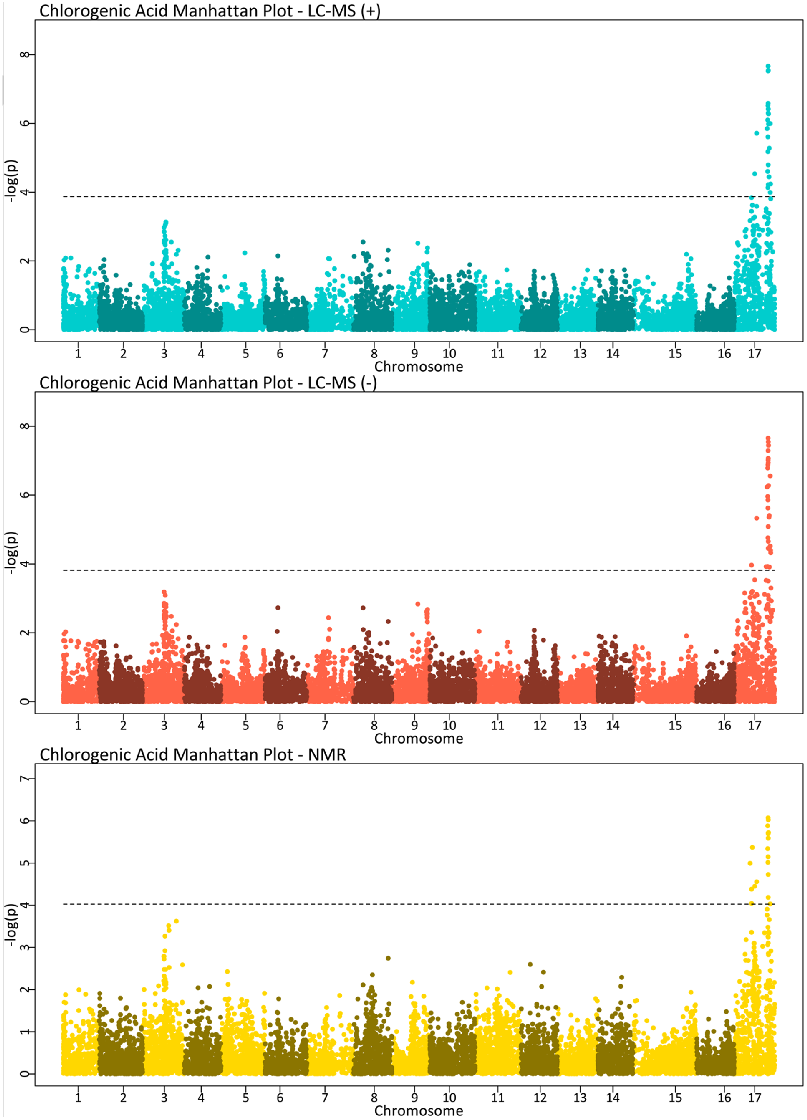
Manhattan plots of chlorogenic acid phenotypic measurements from LCMS (+), (−), and NMR. Alternating colors were used to help delineate neighboring chromosomes. The dashed line indicates an FDR-corrected q-value equivalent to P =.05.

**Fig. 7.**
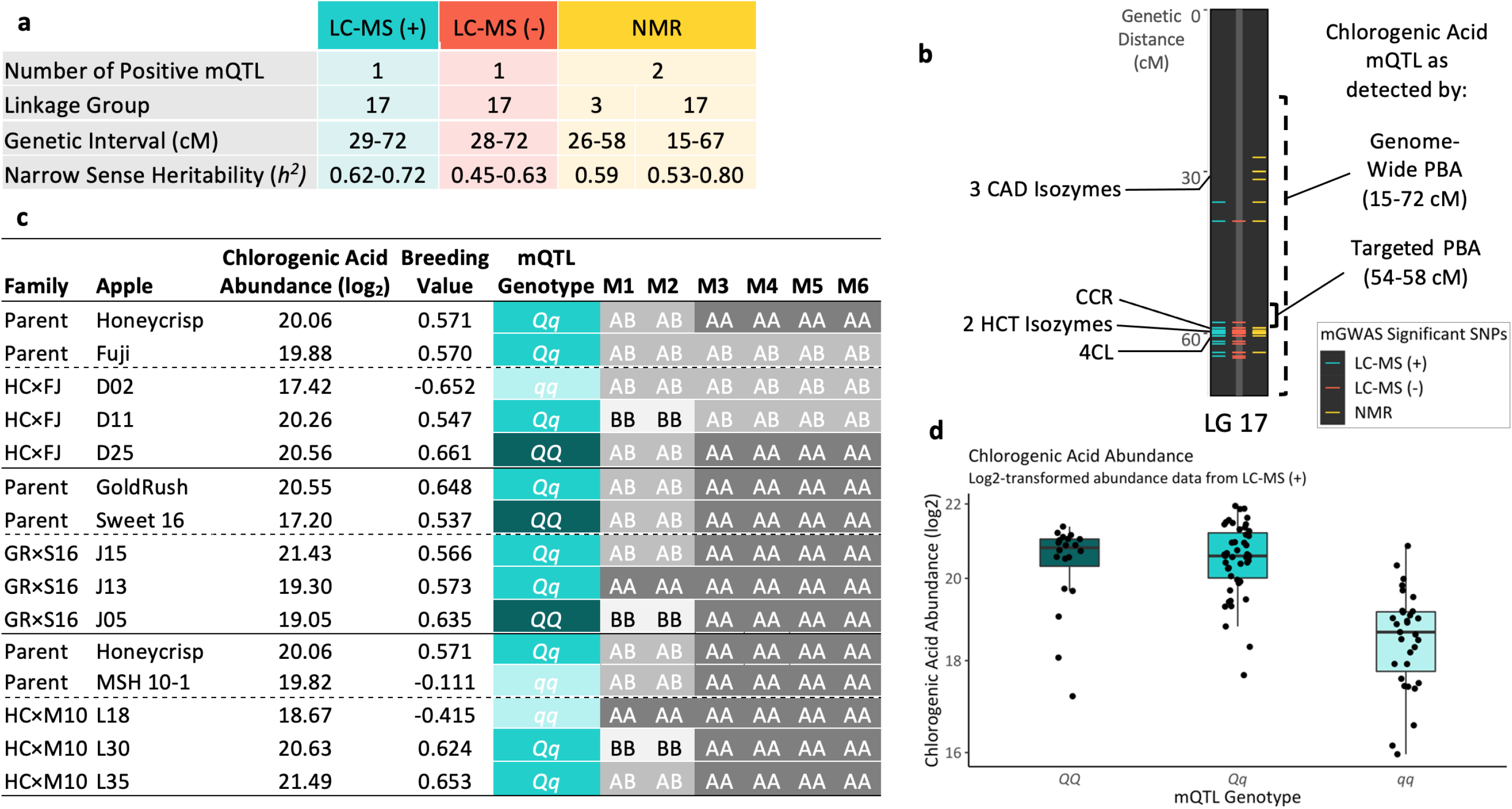
**(a)** Results for the genome-wide pedigree-based analysis (PBA) of chlorogenic acid abundance data for LC-MS (+), (-), and NMR datasets. Number of positive mQTL was determined by a minimum 2ln(Bayes factor) of 2. Genetic interval and narrow sense heritability estimates are recorded in ranges determined from three replicates of the FlexQTL™ runs for each dataset, **(b)** Linkage group (LG) 17 with colored bars indicating SNPs significantly associated with chlorogenic acid in the mGWAS. The dashed line shows the mQTL range (15-72 cM) determined by the genome-wide PBA. The solid line indicates the mQTL region (54-58 cM) detected by the targeted PBA. On the left side of the LG, candidate genes are labeled by their encoded enzymes: three cinnamyl alcohol dehydrogenase (CAD) isozymes, cinnamoyl-CoA reductase (CCR), two hydroxycinnamoyl-CoA shikimate/quinate hydroxycinnamoyl transferase (HCT) isozymes, 4-coumarate-CoA ligase (4CL) (Sup. Table S25). **(c)** Representative individuals from the three families are presented alongside their log2 chlorogenic acid abundance from LC-MS (+), genomic breeding value, mQTL genotype, and haplotype with M1-6 corresponding to markers 15065, 15068, 15076, 15079, 15080, and 15082. “A” corresponds to the most common allele and “B” to the least common allele of the SNP. **(d)** Boxplots show log_2_ chlorogenic acid abundance of individuals (n=124) grouped by their mQTL genotype assigned by the targeted PBA. The deviation from linearity supports dominance gene action in chlorogenic acid abundance of apple fruit.

The findings from the genome-wide PBA in FlexQTL™ are listed in Fig. 7a. Both LC-MS (+) and (-) showed one mQTL for chlorogenic acid. The Bayes factors (2ln(BF)) for replicate FlexQTL™ routines for both MS ionization modes indicated a range of strong (5-10) to decisive (>10) evidence for one mQTL. The results from NMR consistently detected a positive (2-5) mQTL on LG 17 that neared strong evidence for one mQTL. Additionally, a positive signal for LG 3 was detected in one replicate of the NMR routines. In all other PBA runs for chlorogenic acid, the signal on LG 3 fell just short of the positive delineation. The table further outlines a consistent genetic interval for the chlorogenic mQTL on LG 17 from 15-72 cM (Fig. 7a-b). Additionally, narrow sense heritability was high for each metabolomic dataset with an overall range of 0.45-0.80.

The targeted PBA of this long region narrowed the mQTL to a locus of 4 cM (54-58 cM; Fig. 7b) using IBD and posterior probabilities from MCMC. This region fell between markers 15060 and 15086 (Chr17:26,433,748..26,920,509), corresponding to a physical distance of 487 kb at the bottom of LG 17. The probability of the region containing components influencing chlorogenic acid abundance was above 0.61. In that locus, haplotypes were constructed based on polymorphic markers (15065, 15068, 15076, 15079, 15080, and 15082) in the progenies. For each apple variety, genotypes for these markers were assembled alongside the observed abundance for chlorogenic acid measured via LC-MS (+) and two outputs from FlexQTL™: the estimated breeding value (additive + dominance effects) and the most probable mQTL genotype (i.e., *QQ*, *Qq*, *qq*) (Sup. Tables S23-S24). An excerpt is presented here to illustrate the patterns observable between haplotype, chlorogenic acid abundance, and breeding value (Fig. 7c). Fig 7d illustrates the dominance effect for the chlorogenic acid mQTL.

Linkage group 17 houses seven genes encoding four enzymes from the phenylpropanoid pathway: three cinnamyl alcohol dehydrogenase (CAD) isozymes, cinnamoyl-CoA reductase (CCR), two shikimate O-hydroxycinnamoyl-CoA transferase (HCT) isozymes, and 4-coumarate-CoA ligase (Daccord et al., 2017; Jung et al., 2019) (Fig 7b; Sup. Table S25).

Our pipeline demonstrates the feasibility and benefits of multi-omic integration in apple. Genomic and metabolomic datasets were leveraged simultaneously to gain insight into genetic control of metabolite production in fruits. The novel workflow was developed to prioritize putative genotype-phenotype associations characterized by mGWAS for final analysis via genome-wide and targeted PBA to progress towards understanding relationships between genetic components and metabolite abundance.

## Discussion

### Detecting mQTL in Breeding-Relevant Germplasm by Combining mGWAS and PBA

To overcome the limitations of QTL mapping in bi-parental population (Table 1), a PBA approach has been implemented to analyze breeding germplasm (Bink et al., 2014; Fresnedo-Ramírez et al., 2015, 2016; Guan et al., 2015; Cai et al., 2017). The PBA approach analyzes several pedigree-connected families, taking advantage of IBD (Bink et al., 2014), a principle based on knowledge of haplotype inheritance over generations. It evaluates alleles of recent progenies based on alleles of founding cultivars (van de Weg et al., 2004). Important breeding parents and progeny make up the pedigree-related families, so QTL are evaluated in a more breeding-relevant context than in bi-parental populations. This allows evaluation of individuals from a variety of genetic and even environmental backgrounds because samples are taken from existing breeding programs.

**Table 1.** Comparison of genotype-metabolite integration studies in apple. If a breeding-relevant collection of apples was used, the analysis type is indicated as a metabolite genome-wide association study (mGWAS) or pedigree-based analysis (PBA).

| Study | Population Type |  | Phytochemical Analysis |  |
| --- | --- | --- | --- | --- |
|  | Bi-Parental | Breeding | Targeted | Untargeted |
| Khan et al., 2012. *J Exp Bot.* | × |  |  | LC-MS (-) |
| Chagné et al., 2012. *BMC Plant Biol.* | × |  | Polyphenols |  |
| Verdu et al., 2014. *PLoS ONE.* | × |  | Polyphenols |  |
| Guan et al., 2015. *Mol Breeding.* |  | PBA | Sugars |  |
| Gutierrez et al., 2018. *Tree Genet Genomes.* | × |  | Dihydrochalcones |  |
| McClure et al., 2019. *Hortic Res.* |  | mGWAS | Polyphenol |  |
| Christeller et al., 2019. *Sci Rep.* | × |  | Pentacyclic<br>Triterpenes |  |
| Bilbrey et al., 2021. Reported here. |  | mGWAS<br>& PBA |  | LC-MS (+/-)<br>& NMR |

PBA would be the most powerful method for mQTL detection in the pedigree-connected population studied here; however, due to the intensive nature of the routines, the 10,000 metabolomic features could not all be realistically analyzed via this method with the current tools available and the Bayesian approach used by FlexQTL™. Thus, novel, high-throughput prioritization schemes to determine features of interest were developed to bridge the gap between metabolomic analysis and PBA. The scheme was based on less computationally intensive mGWAS applied across the nested population sets (Progeny, Pedigree, Diverse), which could act as a filter, selecting the most important metabolites for analysis using PBA.

Our rationale for dividing the germplasm into three sets was adopted based on the assumption that genetic segregation of metabolite production should have clearest patterns in the progenies. In the mGWAS analyses, the addition of some accessions that are phenotypic outliers, without any consideration of their relationship to the breeding program, would likely lead to detection of spurious genotype-phenotype relationships (Alvarez Prado et al., 2019). Therefore, marker-metabolite associations that retain signal across all three iterations of the analysis should be robust across a wide variety of apple germplasm, as represented by the individuals in this study. The advantage of this approach is generalized applicability of marker-metabolite associations in a wide range of breeding germplasm. Furthermore, this approach reduces false positive SNP-feature associations, as detection across each population is required to remain in the analysis.

While mGWAS alone is useful to understand the genomic architecture for a phenotype (Korte Farlow, 2013), it does not capture the inheritance of each allele through a pedigree despite utilization of a kinship matrix and principal components to correct models. Additionally, our mGWAS analyses included individuals outside the three pedigree-connected families, so the pedigree relationships were only relevant to the portion of the analyses involving individuals related to the breeding progenies. Thus, for estimation of genetic parameters, such as variance explained by an mQTL for a specific phytochemical, and to track the IBD probability of alleles across the pedigree, the Bayesian PBA in FlexQTL™ provides additional information by generating values for additive and dominance effects. It also enables the construction of haplotypes for further exploration of the polymorphisms in the mQTL regions to narrow down candidate markers for marker-assisted selection.

### Benefits of Omics Integration and Using Multiple Metabolomics Approaches

The composite mQTL chromosome map (Fig. 5) displays both genomic areas of interest and SNPs associated across the metabolomic datasets, demonstrating the advantage of applying high-resolution MS and NMR together. This diagram is the first of its kind. Loci with signal across all three metabolomic approaches provide additional confidence in mQTL. Three significant associations also provide chemical information – that those metabolites are observable via both MS ionization modes as well as via NMR. These data suggest that metabolites significant across all three datasets exist in sufficient concentration (>1 μmol/L) to be observed via NMR, which is the least sensitive approach. For example, at the bottom of LG 17, mQTL that are associated with features across the three datasets are visible. The location of the NMR bin (i.e., chemical shift) provides additional structural information about the metabolite expressing that peak. This enables the narrowing down of potential metabolite classes, as shown in Sup. Fig. S8, and limits the number of theoretical formulas matching the accurate mass, aiding in the metabolite identification process. This is particularly valuable as metabolite identification is the major bottleneck of metabolomics studies (da Silva et al., 2015).

Conversely, at the tops of LGs 2 and 3, there are many significant mQTL found across the two MS datasets. It is likely that these features are present in concentrations too low (<1 μmol/L) to be well-characterized by NMR. The presence in one LC-MS ionization mode (and absence in the other) can also provide useful structural information, as certain compounds are more or less likely to gain or lose a proton. Our finding of more significant features from the LC-MS (-) prioritization than from LC-MS (+), despite the positive mode dataset having more features, is consistent with increased analyte ionization in positive mode and increased sensitivity in negative mode (Antignac et al., 2005). In the case of apple, this difference might be explained by the fact that flavonoids, a diverse class of compounds, tend to ionize better in negative mode than positive (López-Fernández et al., 2020). If the chosen apple varieties represent a high diversity of flavonoids, this could account for more significant features passing the thresholds for LC-MS (-). Similarly, areas displaying significance for NMR data only could indicate an area controlling abundant compounds that do not ionize well using MS or are not well-retained on the column in reversed-phase LC and are therefore preferentially analyzed by NMR, such as sugars and other very polar analytes. Though NMR is an inherently less sensitive approach than MS, our work demonstrates the complementarity of using both techniques.

These observations were important clues to strengthen confidence in understanding the classes of compounds being controlled by certain areas of the apple genome. A great advantage of this approach was also to see that parallel results in mQTL detection were evident across mGWAS analyses. This increases confidence in detection of truly significant SNP-feature associations as opposed to chance results from multiple comparisons. These conclusions further strengthen the utility multi-omic integration provides toward additional leverage for compound identification, as well as confidence in results.

### mQTL Hotspot on Linkage Group 16

The mQTL hotspot on the top of LG 16 is consistent with results from mQTL mapping studies by Khan, Chibon, et al. (2012) and Chagné, Krieger, et al. (2012) as well as the mGWAS in McClure et al. (2019). This consistency in findings confers confidence in the validity of our approach for mQTL detection using untargeted metabolomics data in a breeding-relevant population. Without complete compound identification, we can infer that this hotspot contains genetic elements that exert control over production of polyphenols. This is possible due to the innate characteristic of NMR that chemical shift (ppm) is determined by compound structure and therefore chemical class. The NMR bins associated with the SNPs in the hotspot region are located from 8-6 ppm (Sup. Fig. S8), indicating that they represent compounds that contain aromatic hydrogens, likely phenolics and aromatic amino acids (Eisenmann et al. 2016; Iaccarino et al. 2019).

### mQTL Hotspot on Linkage Group 17 including Chlorogenic Acid

Chromosome 17 has not been noted as particularly remarkable in previous apple genomics studies, so an over-representation of mQTL on this linkage group led us to investigate further. The LC-MS features, (+) 97 and (-) 131, significantly associated with SNPs on LG 17 were cross-referenced with features from data-dependent MS/MS that were identified by spectral library matches in GNPS. The comparisons for both ionization modes yielded chlorogenic acid as one such match. Furthermore, five of the seven NMR bins (7.1-7.09, 6.33-6.32, 2.17-2.16, 2.15-2.14, and 2.14-2.13 ppm) with putative mQTL on LG 17 showed chemical shift distribution and multiplicity expected for chlorogenic acid as detailed in HMDB (Wishart et al., 2018). With consistency across metabolomic datasets, as visualized by Manhattan plots (Fig. 6) for these LC-MS and NMR features, we chose chlorogenic acid to be further evaluated via PBA for mQTL validation. Furthermore, chlorogenic acid was selected because its consumption is associated with decreased risk of chronic health concerns such as: diabetes, cancer, inflammation, and obesity (Tajik et al., 2017).

Previous studies have identified an mQTL on the bottom of LG 17 for chlorogenic acid (Chagné et al., 2012; Khan et al., 2012; Verdu et al., 2014; McClure et al., 2019). Three of these studies were based on bi-parental mapping populations (Chagné et al., 2012; Khan et al., 2012; Verdu et al., 2014). A strong but non-significant signal was detected for chlorogenic acid in a breeding-relevant population using mGWAS (McClure et al., 2019). The stronger mQTL signal we observed could be due to the specific breeding-relevant germplasm used for this study. Another reason for more sensitive detection of mQTL in our mGWAS may be attributed to the core of pedigree-connected individuals chosen for the study that enabled an informative kinship matrix to be included in the model. This suggests, although it is important to use breeding-relevant germplasm in mQTL analysis, having sets of progenies and their pedigree-related individuals strengthens the capability of detection surpassing that of a bi-parental mapping population.

To contextualize the presence of mQTL, genomic recharacterization is needed, but existing knowledge of genes and their encoded enzymes associated with pertinent biochemical pathways can be investigated. The mQTL region detected by the genome-wide PBA along the majority of LG 17 was found to contain seven such genes encoding four enzymes: three CAD isozymes, CCR, two HCT isozymes, and 4CL (Fig. 7b). Of these, the *HCT* gene was also hypothesized to be a candidate gene for the chlorogenic acid mQTL previously identified in two mapping populations: ‘Royal Gala’ × ‘Braeburn’ and a cross of two French cider apples (Chagné et al., 2012; Verdu et al., 2014). This enzyme acts in the phenylpropanoid pathway to transfer quinic acid and shikimic acid to and from molecules upstream of chlorogenic acid (Clifford et al., 2017). However, the other genes have not been previously noted as candidate genes for any chlorogenic acid mQTL in apple. Down-regulation of CAD, *4CL,* and *CCR* results in reduced lignin content with simultaneous increased abundance of upstream metabolites, such as chlorogenic acid (Anterola Lewis, 2002). Thus, these genes are highly likely to be important to chlorogenic accumulation in apple fruit. Furthermore, after conducting the targeted PBA, the smaller mQTL region (Chr17:26,433,748..26,920,509) was 159 kb from *CCR*, which encodes first committed enzyme of the monolignol biosynthetic pathway, a branch off of the general phenylpropanoid pathway (Lacombe et al., 1997). Specifically, down-regulating *CCR* in tomato (*Solanum lycopersicum* L.) decreased lignin biosynthesis, leaving more coumaroyl-CoA esters for use in phenolic compound synthesis (van der Rest et al., 2006). This proof-of-concept example using chlorogenic acid allowed us to uncover previously known relationships, like HCT, as well as discover new gene-metabolite relationships, such as with *CCR*, *CAD*, and *4CL*, in apple.

The same investigative process can be applied to elucidate genetic control of additional compounds. The smaller number of significantly associated features in the NMR dataset indicates that the bulk of the compounds with mQTL for LG 17 are not abundant enough to be detected via NMR metabolomics or mQTL detection is hampered by the nature of binning in NMR, where signal is diluted and distributed across multiple chemical shifts. Therefore, additional scrutiny of our significant features in LC-MS and cross-referencing with GNPS spectral matches may yield more features with LG 17 mQTL that are primed for identification and further validation via PBA. The majority of other features with putative mQTL on LG 17 co-located with the chlorogenic acid mQTL identified via mGWAS (Fig. S7). These findings suggested that the cluster of mQTL detected near the bottom of LG 17 might be metabolically related to chlorogenic acid. This leads to additional hypotheses for the putative identification of other polyphenols with apparent mQTL, such as caffeic acid, co-located at this same locus. This hypothesis is supported by the previously identified mQTL for quercetin-3-O-rutinoside along the entirety of LG 17 (Chagné et al., 2012) and flavonols on the lower half of 17 (Verdu et al., 2014).

### Conclusions and Future Prospects

This study provides a pipeline for high-throughput processing and integration of genomic and metabolomic datasets in diverse, breeding-relevant apple germplasm. The genomic and metabolomic variety in the selected apples led to detection of mQTL across the apple genome. The use of three parallel metabolomic approaches offered complementary interpretation of results that increased confidence in compound identification and detection of significant mQTL through mGWAS and PBA. This workflow enabled detection of 519, 726, and 177 putative mQTL in LC-MS (+), (-), and NMR datasets, respectively. Furthermore, chlorogenic acid represented a proof-of-concept example that this approach was able to characterize and detect consistent mQTL via mGWAS and two-step PBA for metabolites relevant to breeding for quality traits, such as nutrition (Tajik et al., 2017). The mQTL region was interrogated to determine candidate genes related to chlorogenic acid biosynthesis, including previously hypothesized HCT and three genes *(CAD, CCR,* and *4CL)* newly associated with an mQTL for chlorogenic acid production in apple. The data is ripe for additional investigation into feature identities and areas of the apple genome that deserve to be recharacterized for a better understanding of what genes or regulatory elements are present. Subsequently, markers could be developed for mQTL if the compounds are of interest for breeding purposes. Additional study of SNP-feature associations not investigated here could be conducted in the future and likely yield interesting and applicable results. Metabolites or SNPs of interest *a priori* could also be investigated in the data generated in this study. Apple QTL for larger scale phenotypes, such as disease resistance, sweetness, or acidity, could now be compared with mQTL to discern if certain metabolites may contribute to the collective phenotype.

The pipeline could be extended to other tree crops and could be adapted for use with other perennial and annual crops, as well. Platforms that provide feasible assessment of genotype-metabolite relationships are imperative as crop research moves towards metabolome-based increased nutrition, disease resistance, post-harvest quality, and consumer-likability. This is especially important in perennial tree crops, such as apple, which will rely upon marker-assisted breeding to overcome a long breeding cycle and accomplish marked advances to parallel or ideally influence consumer desires and needs.

## Acknowledgements

This study was supported by Foods for Health, a focus area of the Discovery Themes at The Ohio State University and USDA Hatch Funds (OHO01399 and OHO01470), with additional support from USDA SCRI RosBREED (2014-51181-22378). Emma Bilbrey was supported by the University Fellowship from The Ohio State University and the Ohio Agricultural Research and Development Center Director’s Associateship Award. A special thanks for providing apples to the Midwest Apple Improvement Association growers Mitch Lynd, David Hull, and David Doud; Thomas Chao at the USDA Apple Germplasm Repository; The Dawes Arboretum in Newark, OH; and Anna Whipkey at Purdue University. Thanks to Eric van de Weg and HJJ (Herma) Koehorst-van Putten at Wageningen University for consulting and providing data on apple pedigree and PBA analysis inquiries. Likewise, thanks to Mike Sovic at The Ohio State University for assistance in batch script development for analyses in the Ohio Supercomputing Center. Thanks also to Cheri Nemes and Maryssa Starman at The Ohio State University for assistance in sample processing. We appreciate the assistance of Dan Cuthbertson (Agilent Technologies) for guidance in the collection of quality, comprehensive MS/MS spectra. This study made use of the Campus Chemical Instrument Center NMR facility at Ohio State University. The format of our preprint was made beautiful by the HenriquesLab bioRxiv Template in Overleaf.

## Author Contributions

EAB, EH, DM, JFR, and JLC designed the study. EAB, KW, and EH conducted experiments. EAB, JFR, and JLC conducted data analysis. EAB and JLC drafted the manuscript, and all authors read and approved the final manuscript.

## Data Availability

The data that support the findings of this study are openly available in the supplemental information or in online repositories. Raw full scan LC-MS data can be found in MetaboLights at www.ebi.ac.uk/metabolights/MTBLS2327 (Haug et al., 2020). Data-dependent LC-MS/MS raw spectra can be found in the MassIVE database at https://doi.org/doi:10.25345/C5B21P. All dataused in analysis can be found in the Supplementary Information. Code for data processing, analysis, and visualization is available at https://github.com/CooperstoneLab within the “apple” repository.

