## Supplemental Figures and Methods for "Integrating genomics and multi-platform metabolomics enables metabolite QTL detection in breeding-relevant apple germplasm"

The following Supporting Information is available for this article:

**Fig. S1** Apple pedigree chart

**Fig. S2** Boxplots of apple extracts for raw and transformed metabolomics data sets

**Fig. S3** LC-MS PCAs including pooled quality control samples

**Fig. S4** Scree and score plots of SNP data PCAs

**Fig. S5** LC-MS and NMR PCAs without pooled quality control samples.

**Fig. S6** Number of metabolites associated with each SNP in chromosome 16

**Fig. S7** Number of metabolites associated with each SNP in chromosome 17

**Fig. S8** Bins across the NMR spectra with putative mQTL

**Table S1** Sample metadata

**Table S2** mGWAS SNP data

**Table S3** Apple extract drying protocol

**Table S4** mzMine2 parameters for processing LC-MS (+) data

**Table S5** mzMine2 parameters for processing LC-MS (-) data

**Table S6** LC-MS (+) deconvoluted data matrix

**Table S7** LC-MS (-) deconvoluted data matrix

**Table S8** 1D NMR Processed Data Matrix

**Table S9** PBA SNP data

**Table S10** SNP names and metadata

**Table S11** Parentage for A matrix

**Table S12** SNPs for G matrix

**Table S13** LC-MS (+) Diverse -log_10_(P) values from mGWAS

**Table S14** LC-MS (+) Pedigree -log_10_(P) values from mGWAS

**Table S15** LC-MS (+) Progeny -log_10_(P) values from mGWAS

**Table S16** LC-MS (-) Diverse -log_10_(P) values from mGWAS

**Table S17** LC-MS (-) Pedigree -log_10_(P) values from mGWAS

**Table S18** LC-MS (-) Progeny -log_10_(P) values from mGWAS

**Table S19** NMR Diverse -log_10_(P) values from mGWAS

**Table S20** NMR Pedigree -log_10_(P) values from mGWAS

**Table S21** NMR Progeny -log_10_(P) values from mGWAS

**Table S22** Significant metabolomic features from mGWAS

**Table S23** Chlorogenic acid mQTL haplotypes for parents and progenies

**Table S24** Chlorogenic acid mQTL haplotypes for all members of the pedigree

**Table S25** Chlorogenic acid mQTL candidate genes on linkage group 17

**Methods S1** Sample collection and initial processing

**Methods S2** Apple fruit extraction

**Methods S3** Full scan LC-MS method, instrument settings, and data deconvolution

**Methods S4** 1D ^1^H NMR spectroscopy method, instrument settings, and data deconvolution

**Methods S5** Data-dependent LC-MS/MS method, instrument settings, and data processing

**Methods S6** Metabolomics feature identification

**Methods S7** Pedigree-based analysis details

**Fig. S1** Apple varieties and relationships with the three pedigree-connected families.

|  | Both SNP data and metabolomics data collected |
| --- | --- |
|  | Seed Parent |
|  | Pollen Parent |

Jonathan

Rome Beauty

*M. floribunda*

9433-2-2

Golden Delicious

PRI-14-226

Crandall

PRI-669-205

Yellow

Newton

NJ-60837

D1R102T98

PWR37T133

CQR10T17

Illinois #2

Co-op 17

S80ER18T32

Idared

PRI-187-6

Wagener

Red

Delicious

PRI-49-102

Winesap

F2-26830-2-2

9433-2-8

Ben Davis

F2-26829-2-2

Fuji (FJ)

Ralls Janet

Keepsake

MN1627

Honeycrisp (HC)

Northern Spy

Frostbite

GoldRush (GR)

Grimes Golden

MSH 10-1 (M10)

Malinda

Sweet 16 (S16)

HC×M10 Progeny

(n=19)

Duchess of

Oldenberg

GR×S16 Progeny

(n=28)

HC×FJ Progeny

(n=28)

**Fig. S2** Scree plots of the percent variation explained by each principal component (PC) in a principal component analysis (PCA) of SNP calls (n=10,294) used for mGWAS for (**a**) the diverse population (n=124), (**b**) the pedigree subset (n=98), and (**c**) the progeny subset (n=75). The red point indicates the elbow: the last PC included in the mGWAS model (**a**: PC=10, **b**: PC=6, **c**: PC=3). Paired with each population subset is a scores plot inset, in which points are apple genotypes.

**b**

**a**

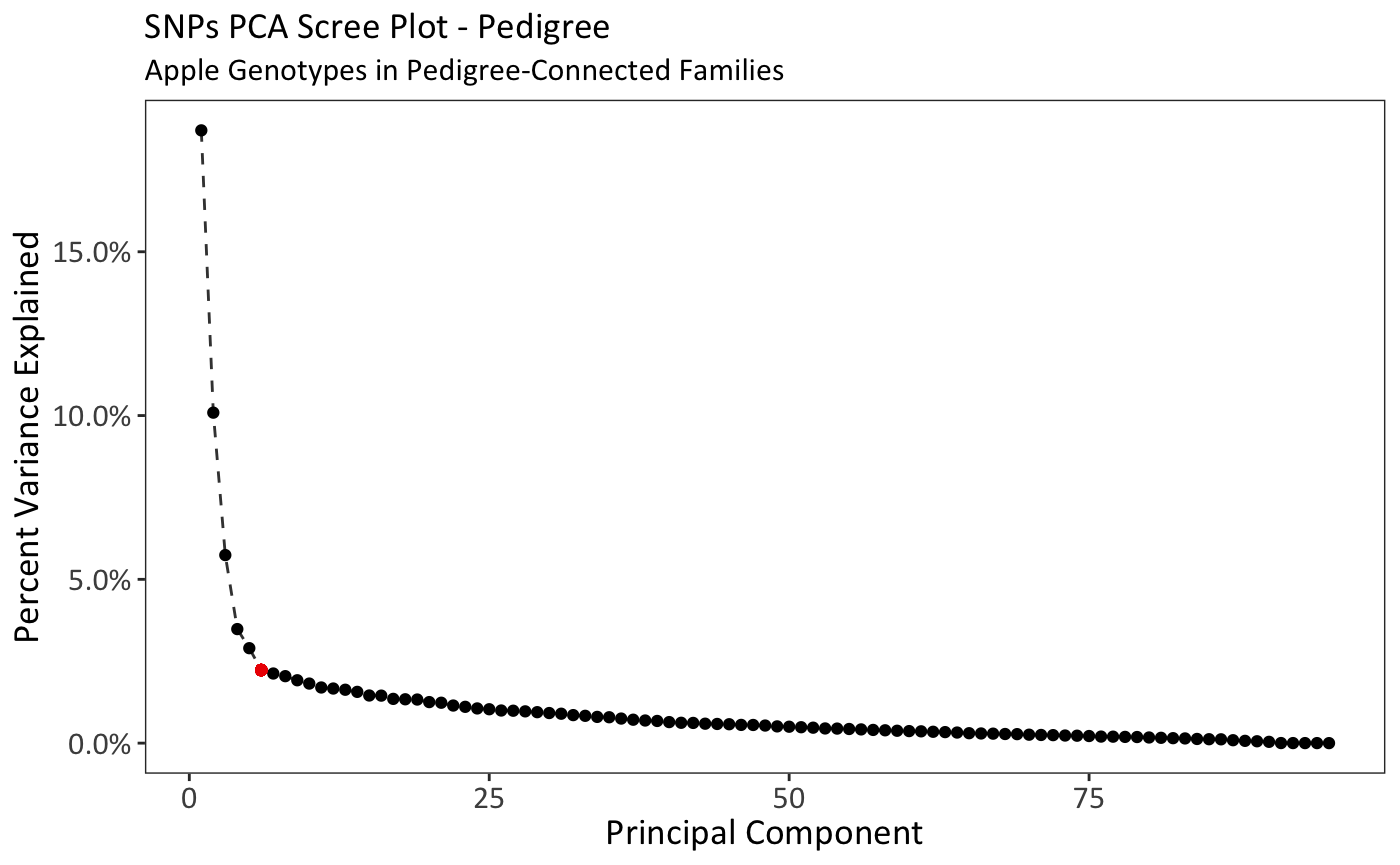

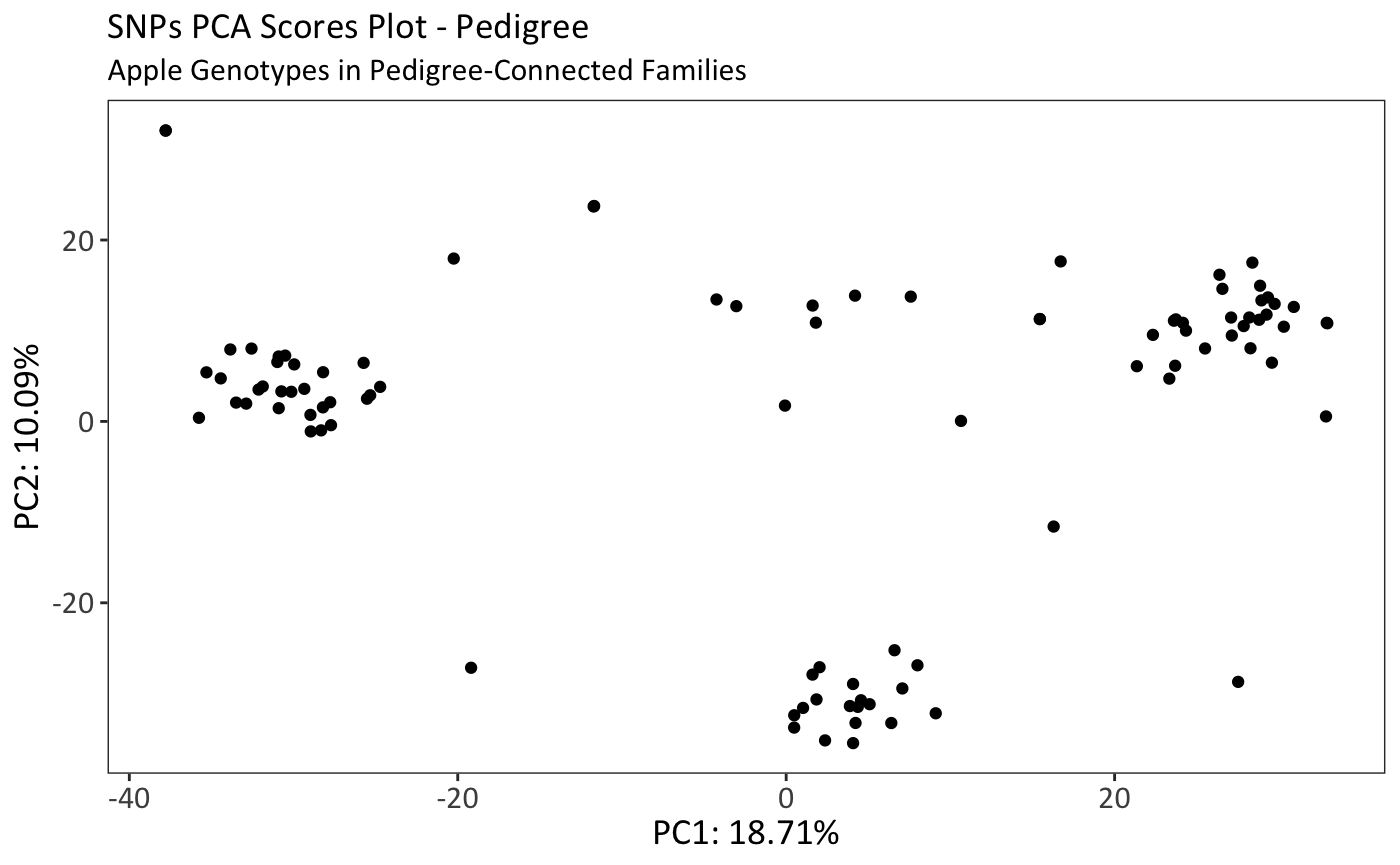

6

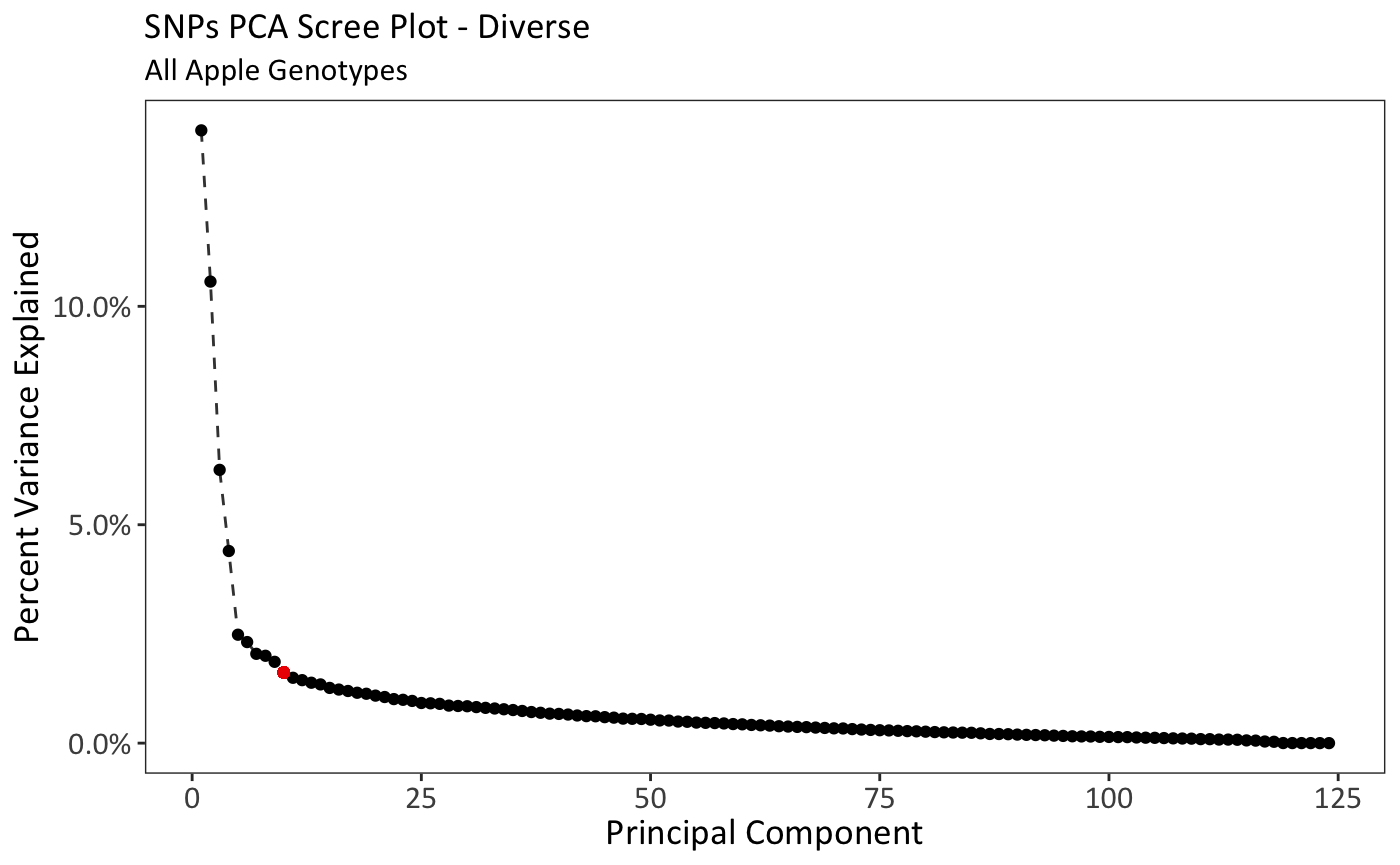

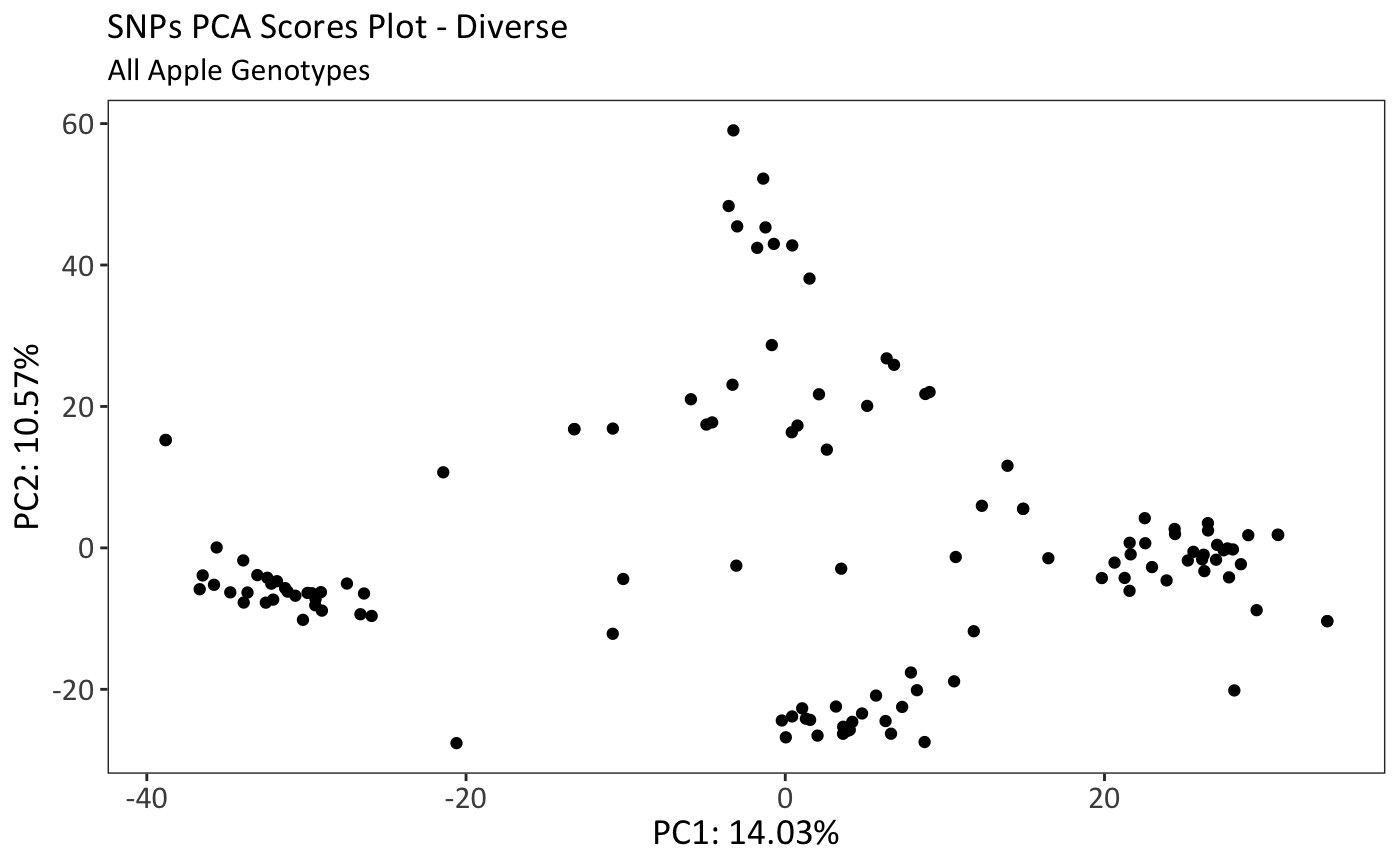

10

**c**

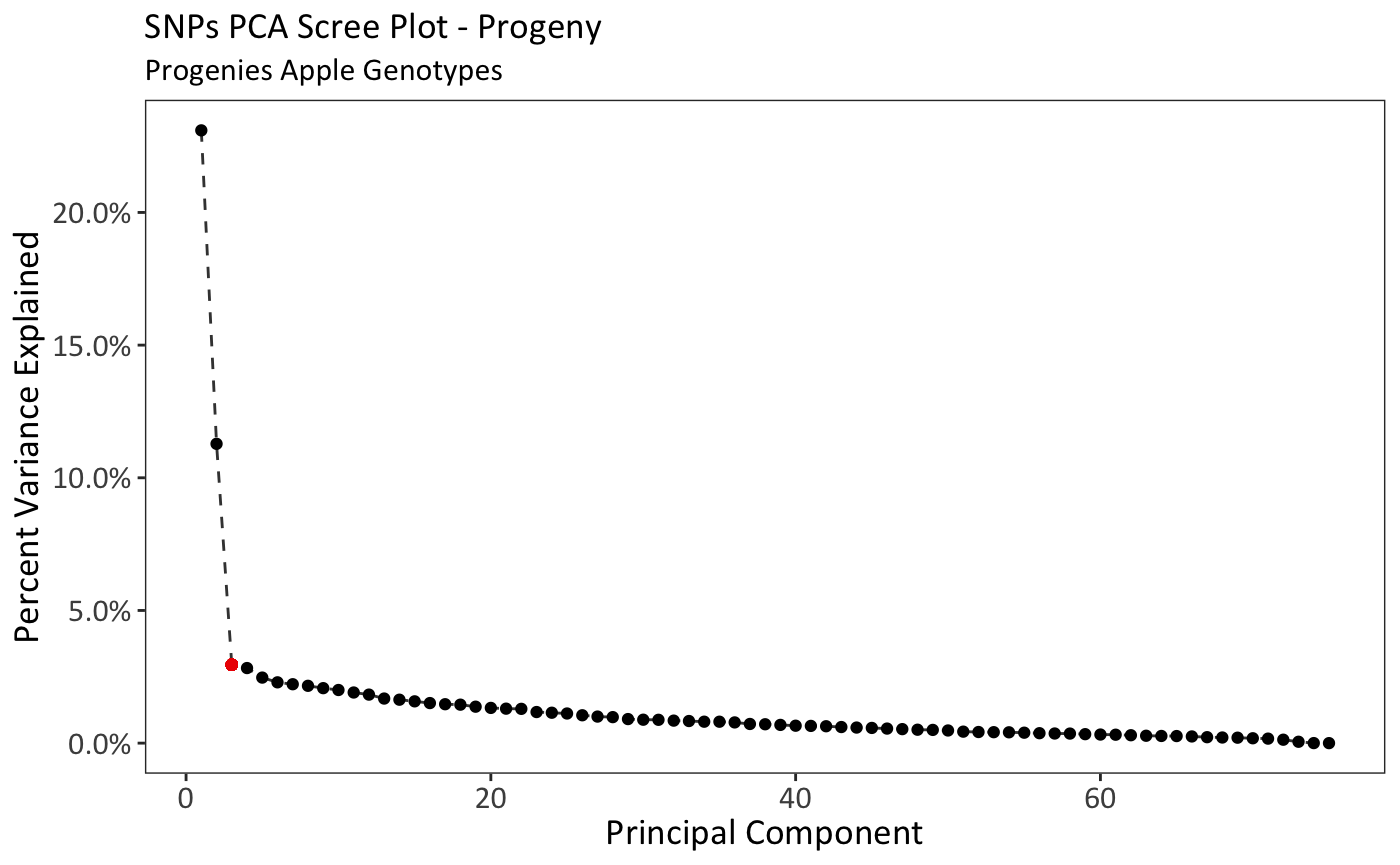

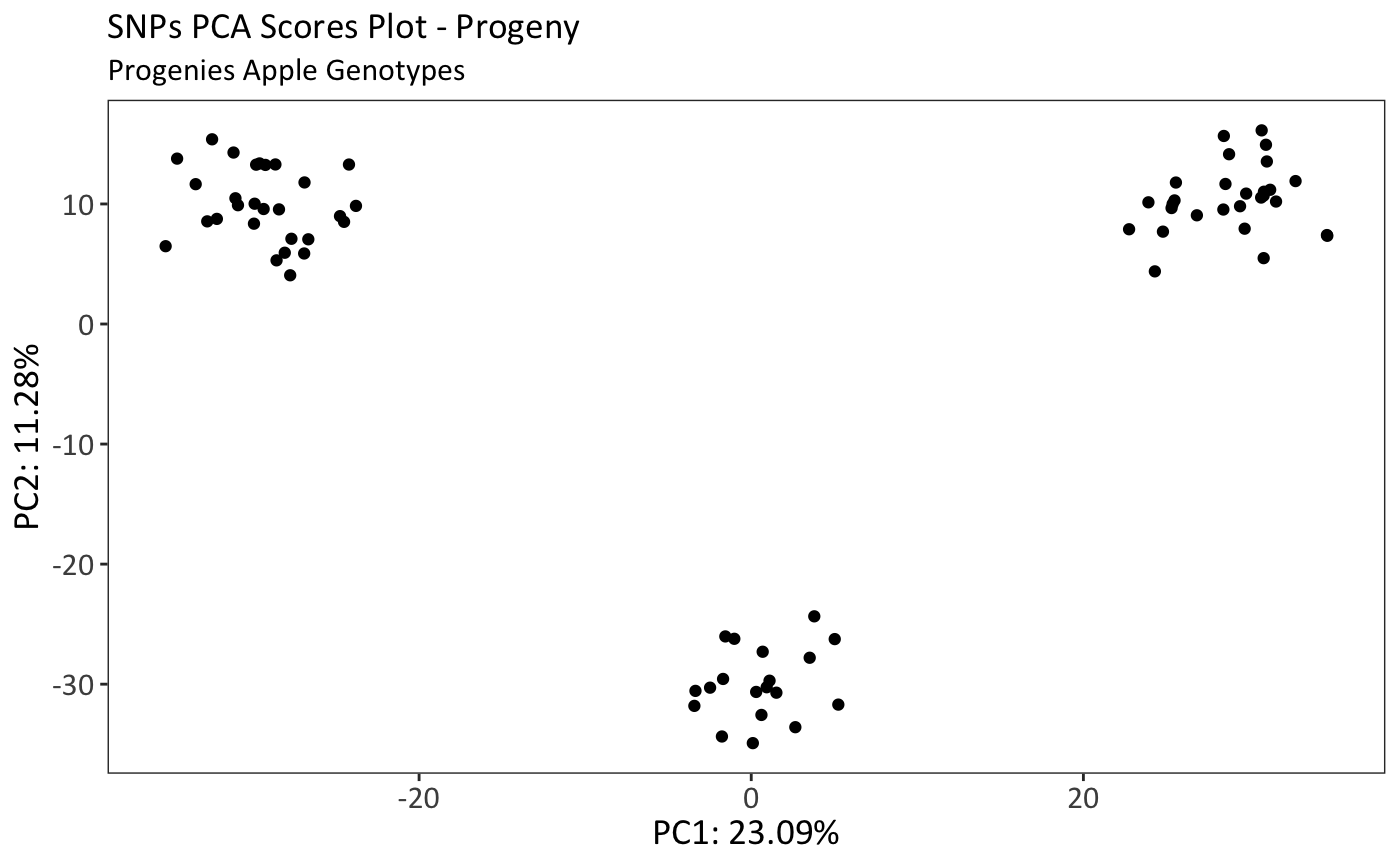

3

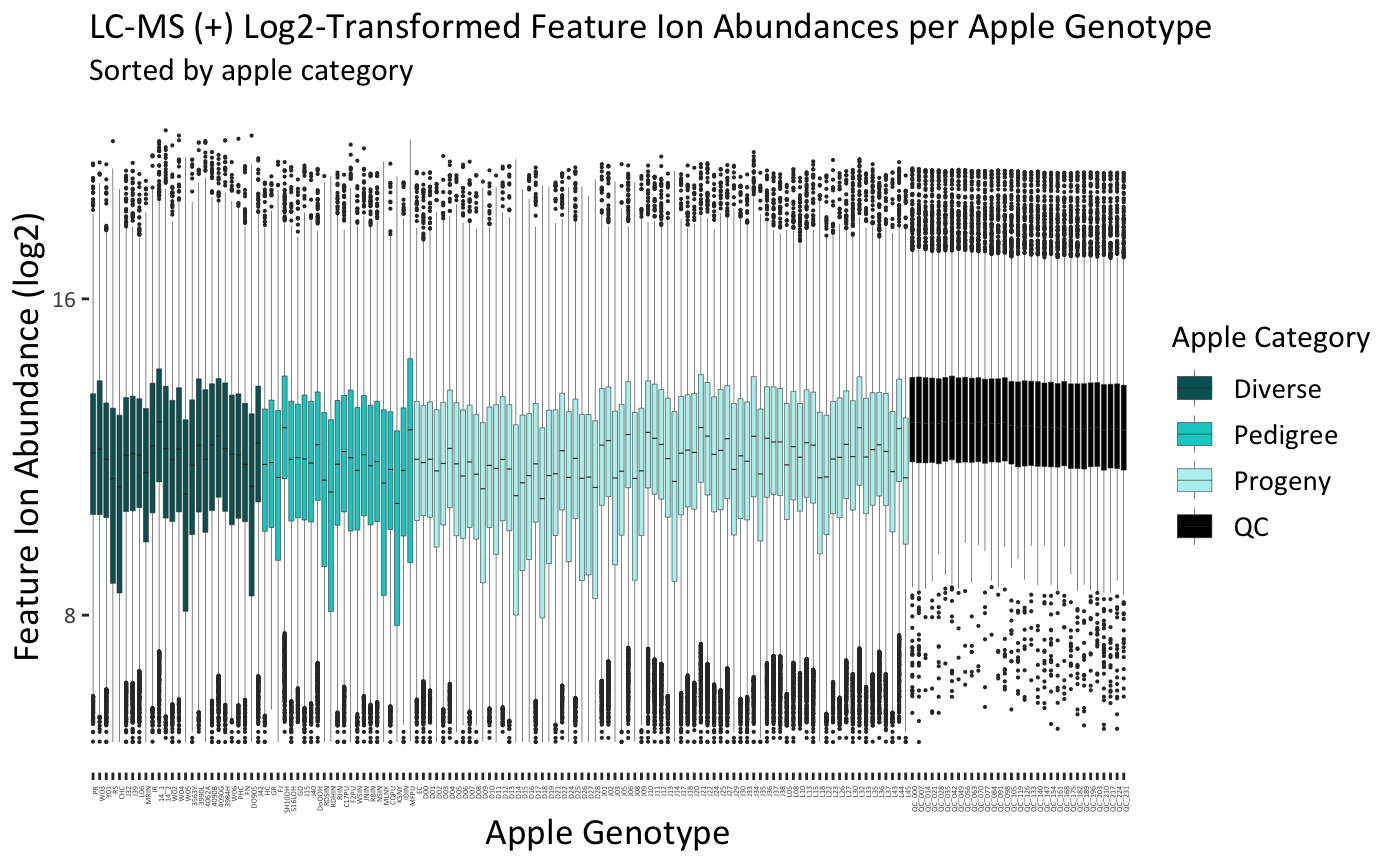

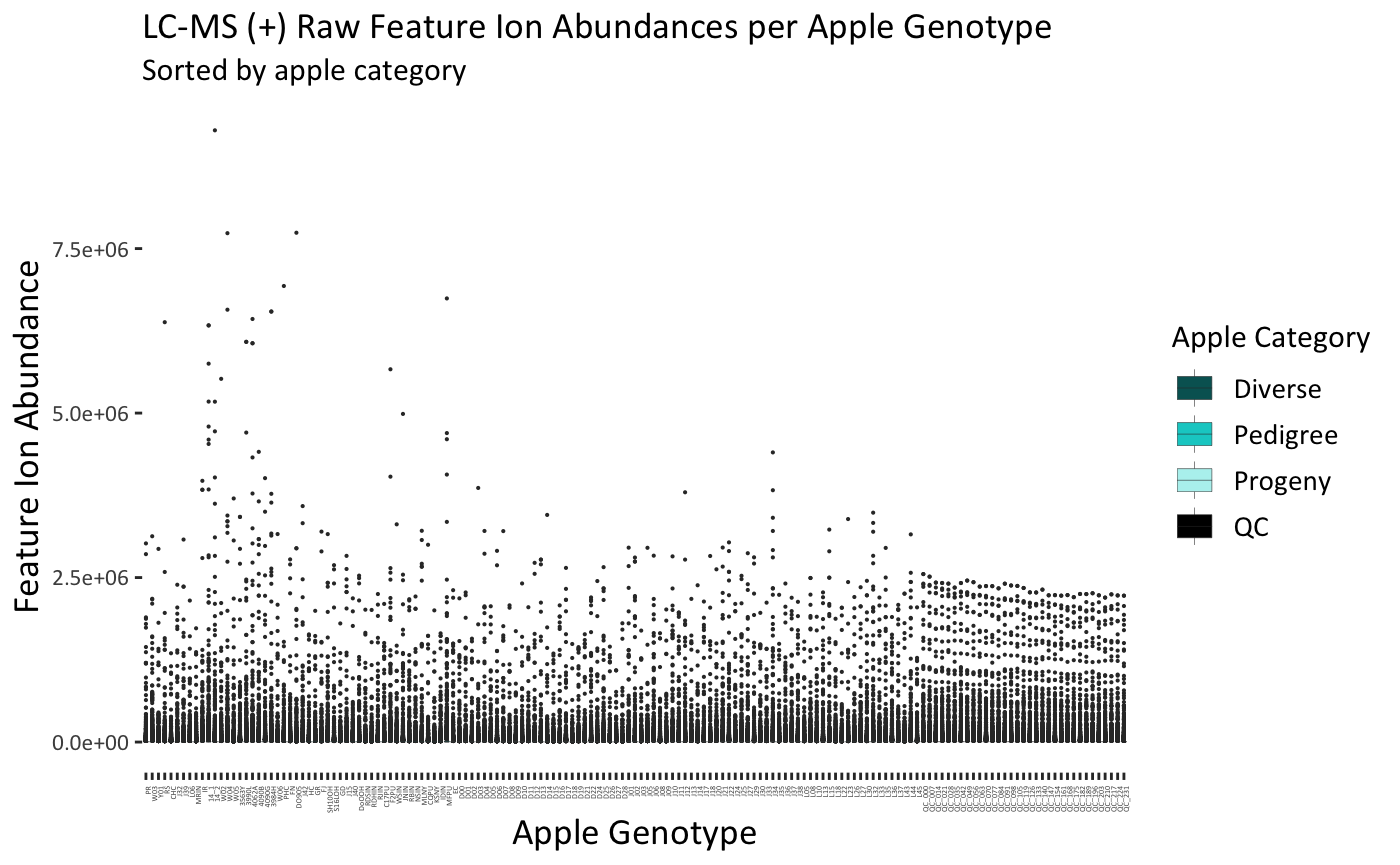
 **Fig. S3** Boxplots for apple extracts analyzed via the three metabolomics platforms: LC-MS (+), (-), and NMR. No samples were identified as notable outliers. Plots for raw and log_2_-transformed data are presented. Missing values were imputed with the lowest value divided by two in the LC-MS datasets.

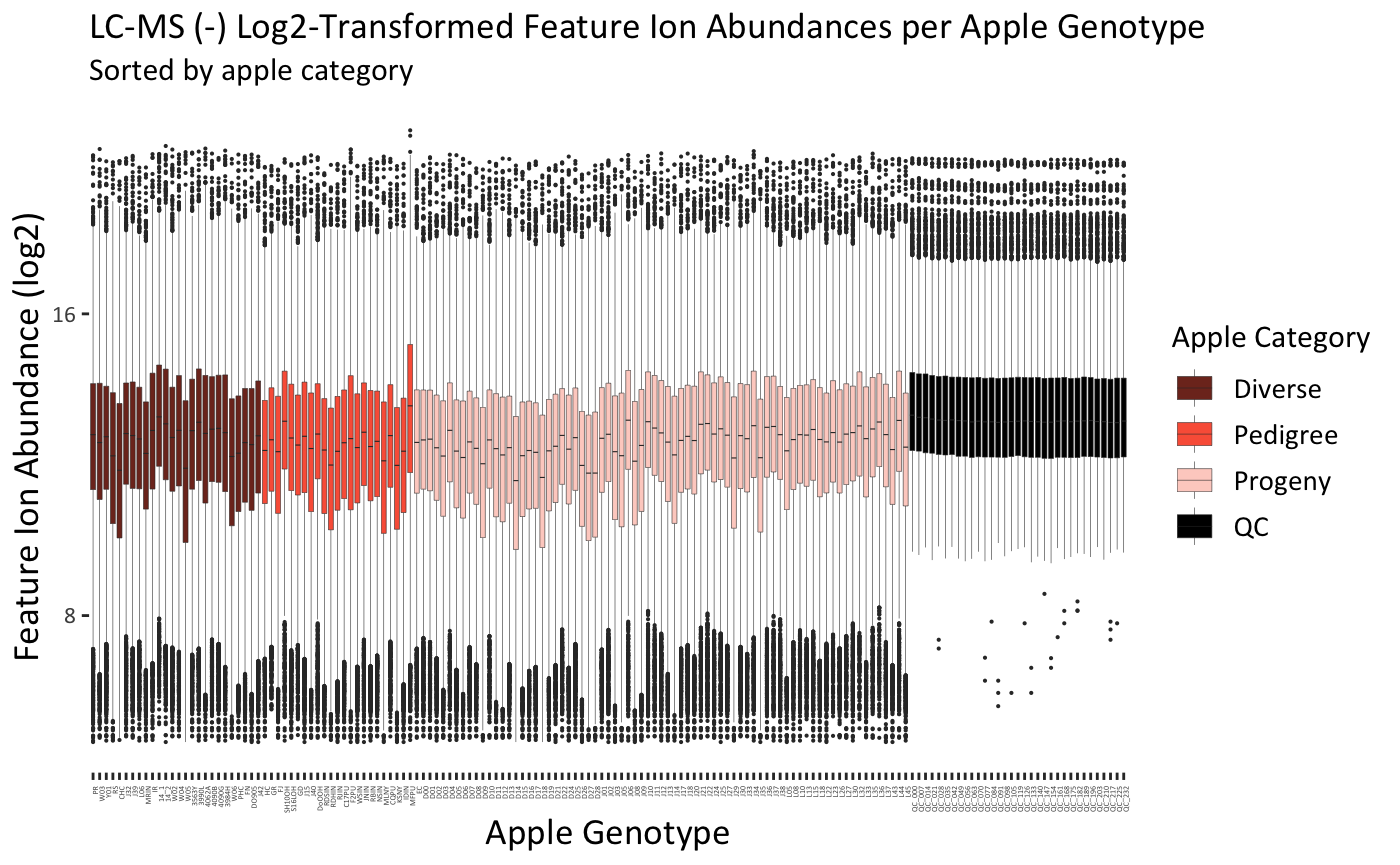

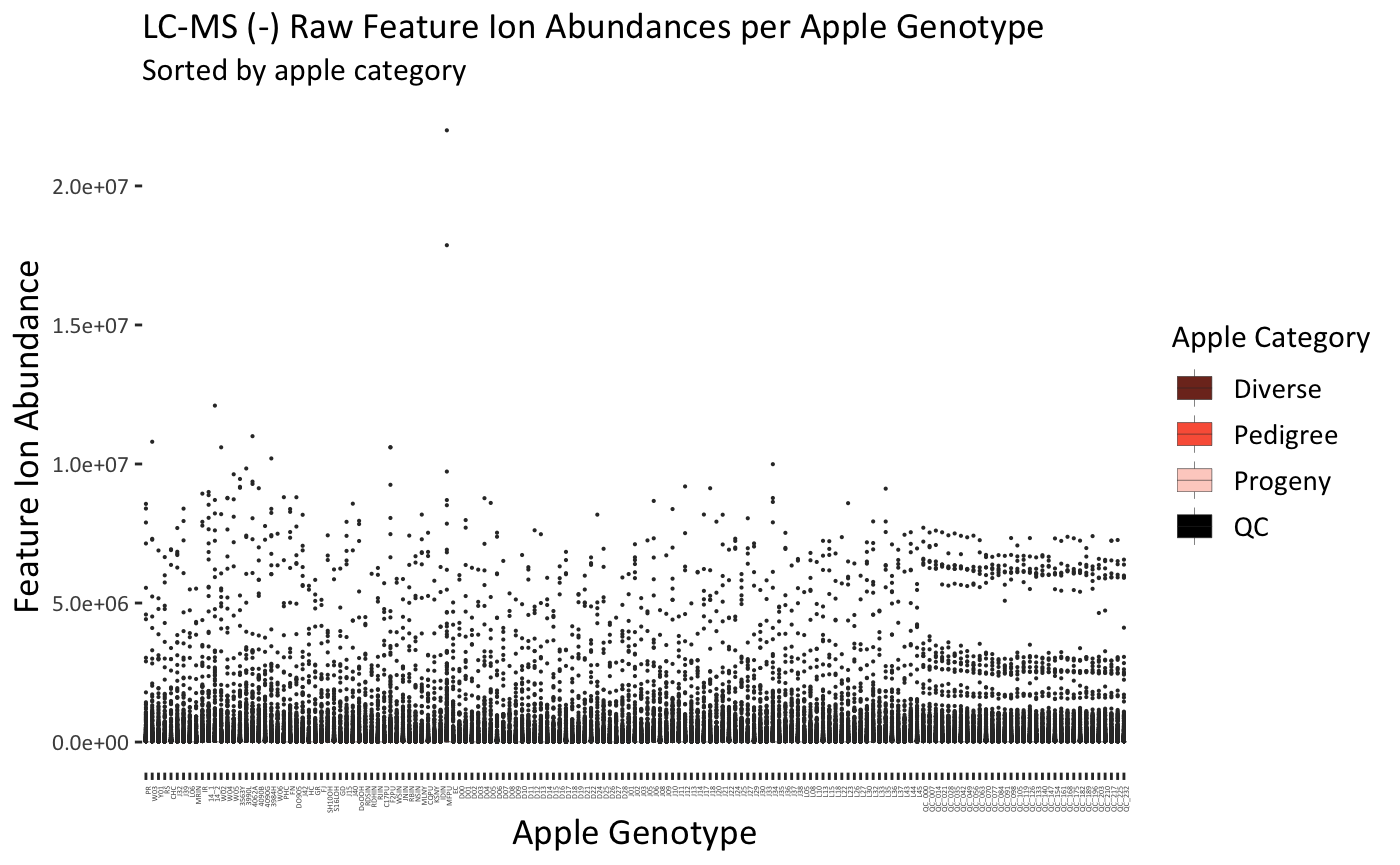

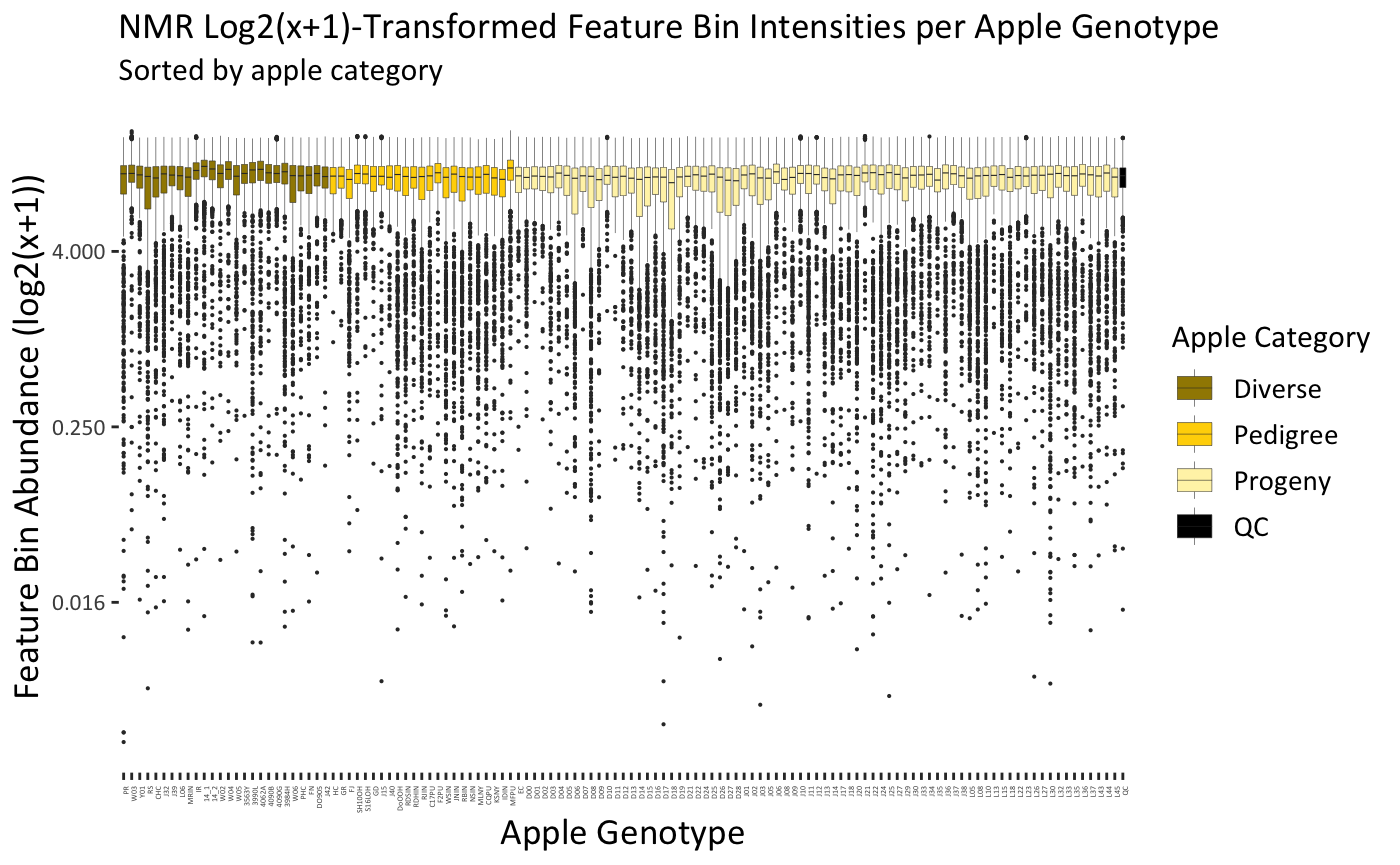
**Fig. S3 cont.** Boxplots for apple extracts analyzed via the three metabolomics platforms: LC-MS (+), (-), and NMR. No samples were identified as notable outliers. Plots for raw and log_2_-transformed data are presented. Missing values were imputed with the lowest value divided by two in LC-MS datasets.

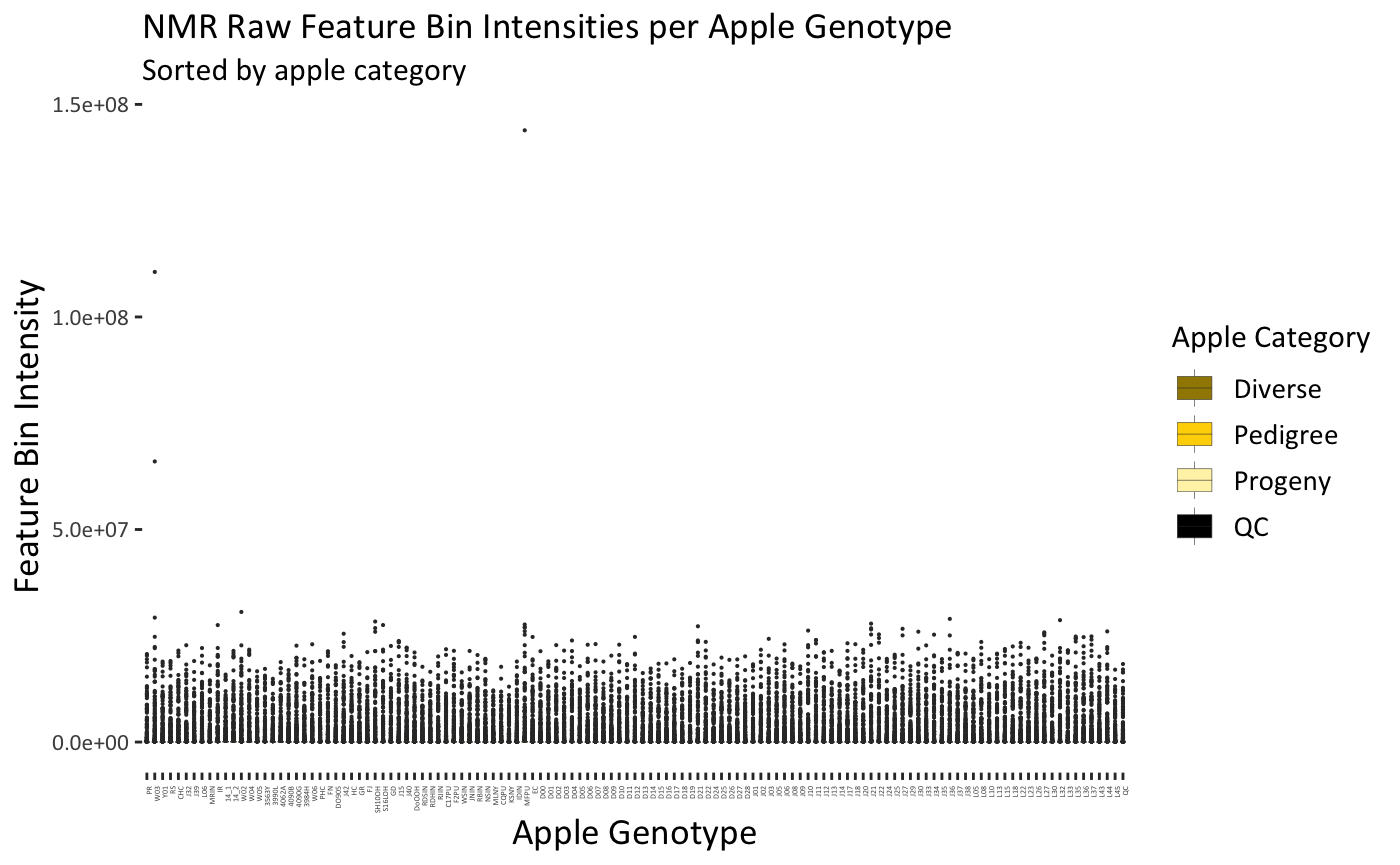

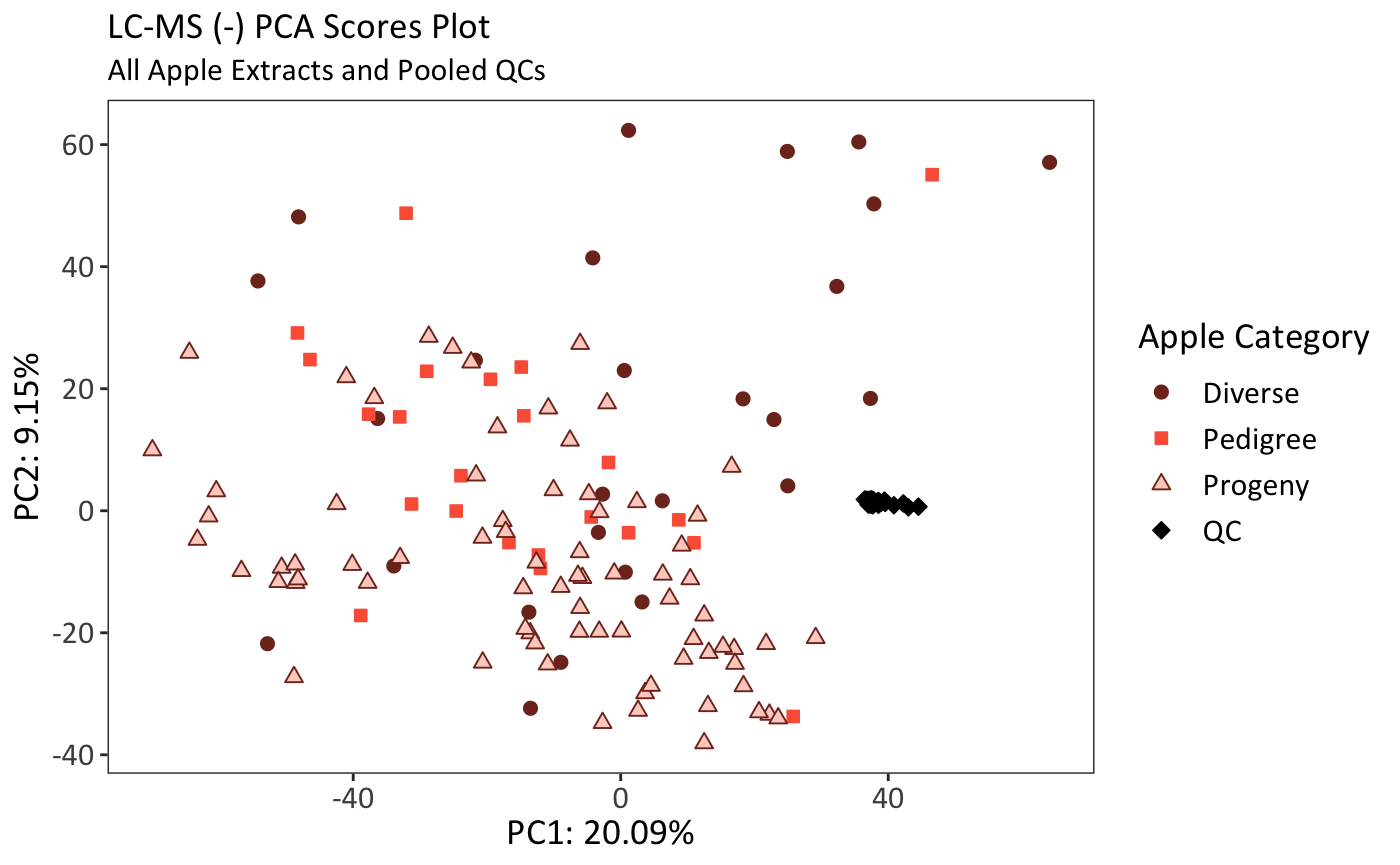

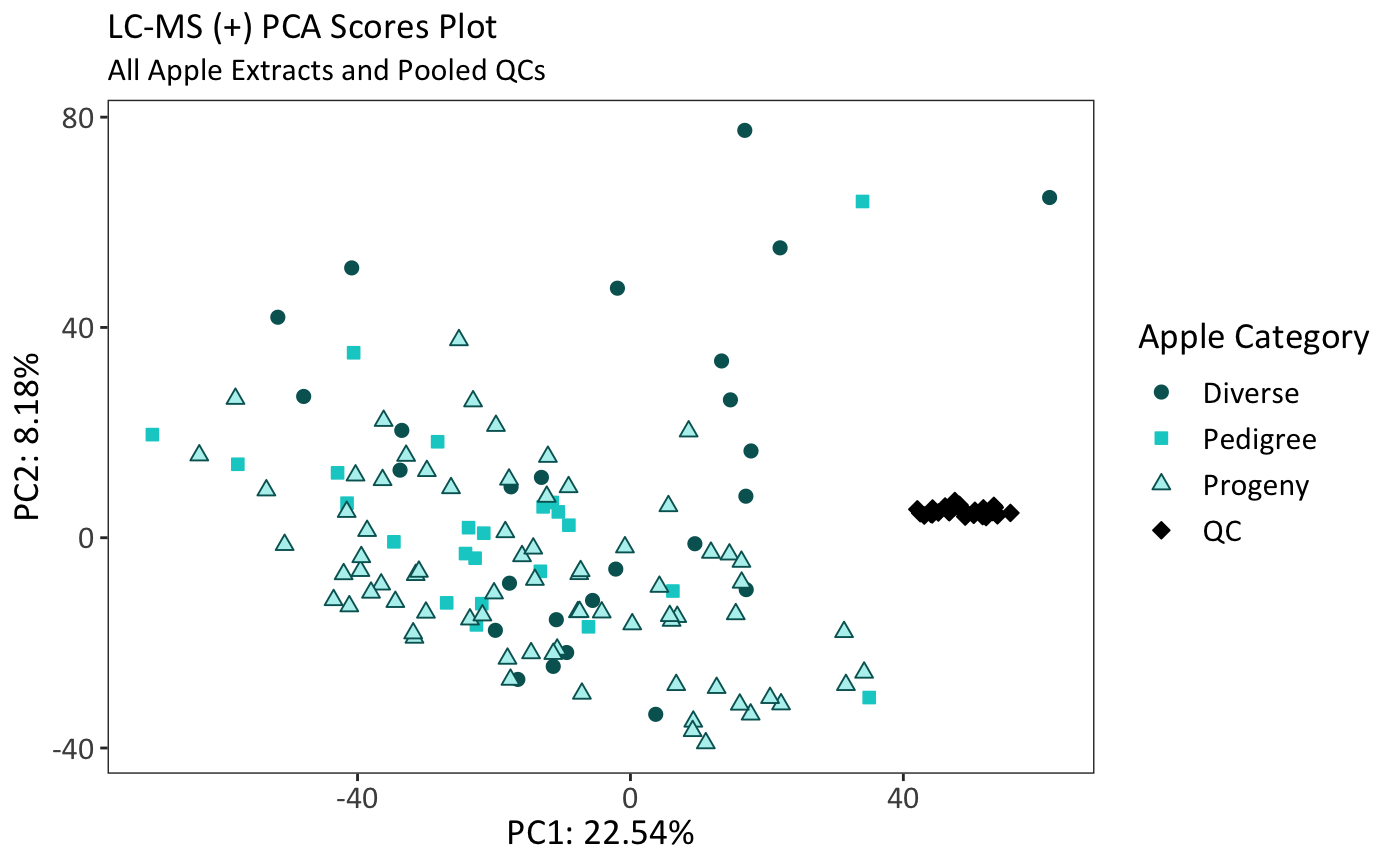
 **Fig. S4** LC-MS (+) and (-) principal components analysis (PCA) of all apple extracts with pooled quality control (QC) samples. QCs clustered tightly, indicating stable data quality in LC-MS analyses. Apple categories are general categories to assess metabolomic diversity in selected samples and do not represent groups being compared in this study. Axes represent the first two principal components and the percent variation explained by each. Data were log_2_-transformed, scaled, and centered.

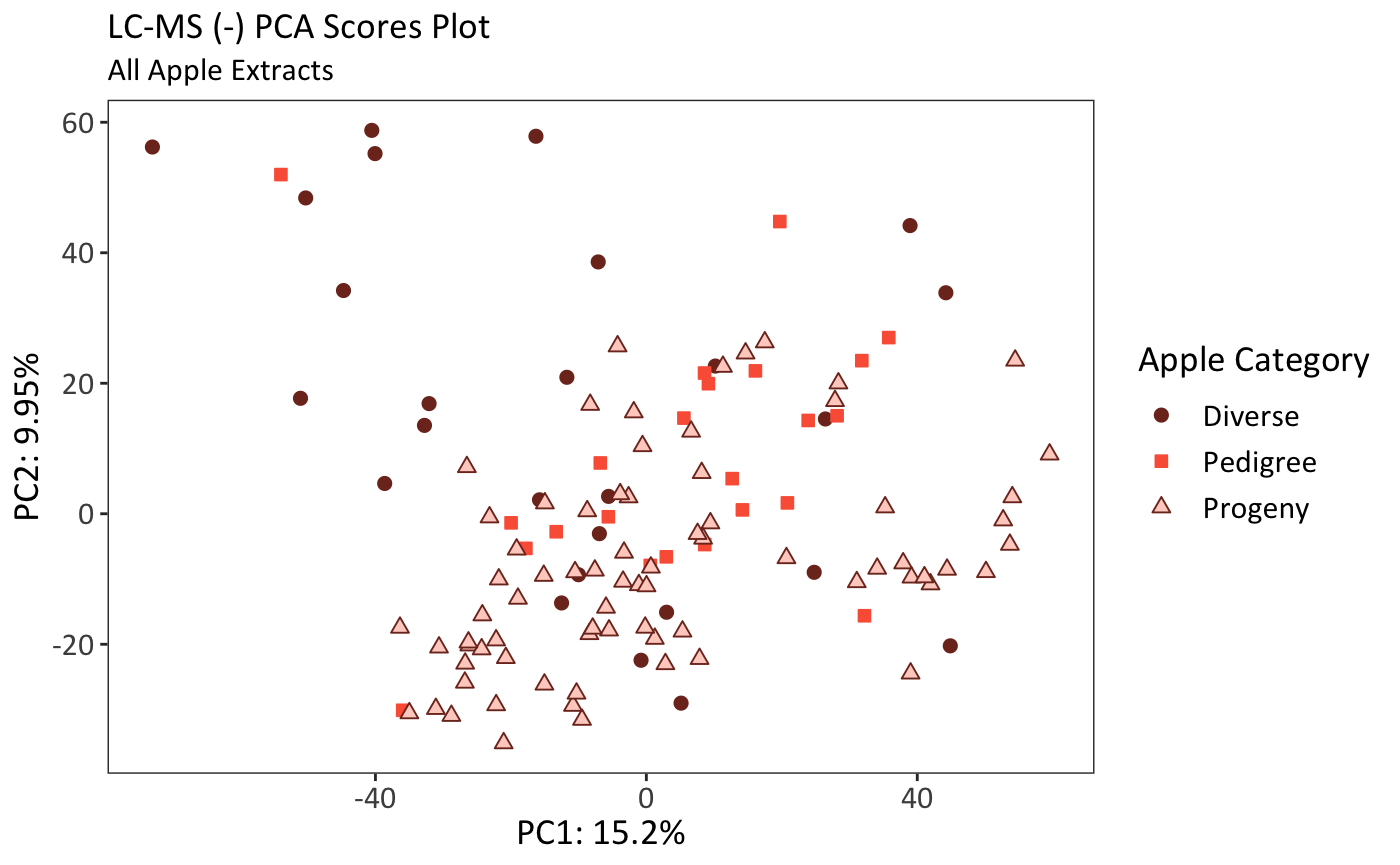

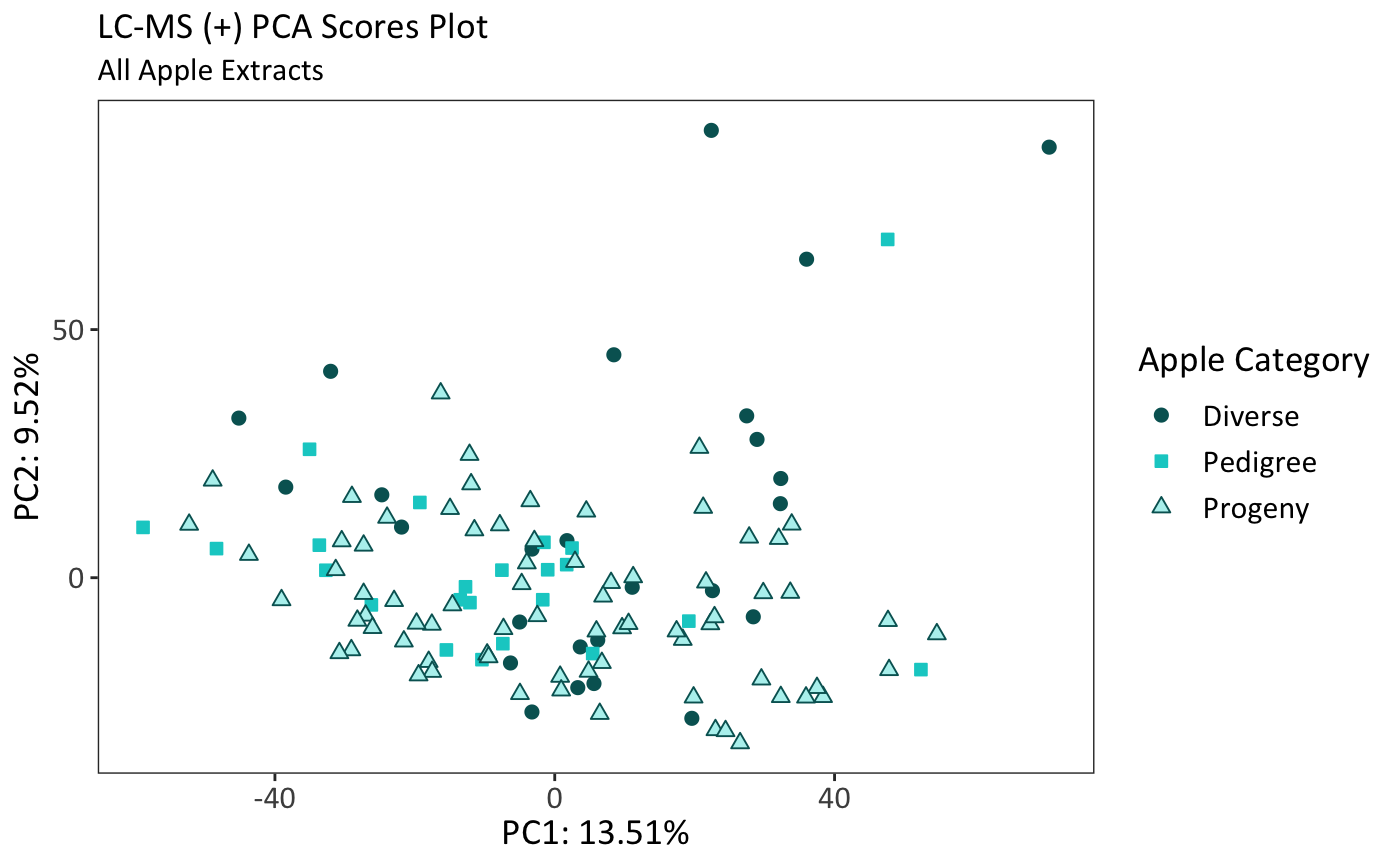
 **Fig. S5** LC-MS (+), LC-MS (-), and NMR principal components analysis (PCA) without pooled quality control (QC) samples. Points are apple genotypes. Apple categories are general categories to assess metabolomic diversity and do not represent groups being compared in this study. Axes represent the first two principal components and the percent variation explained by each. Data was log_2_-tranformed, scaled, and centered.

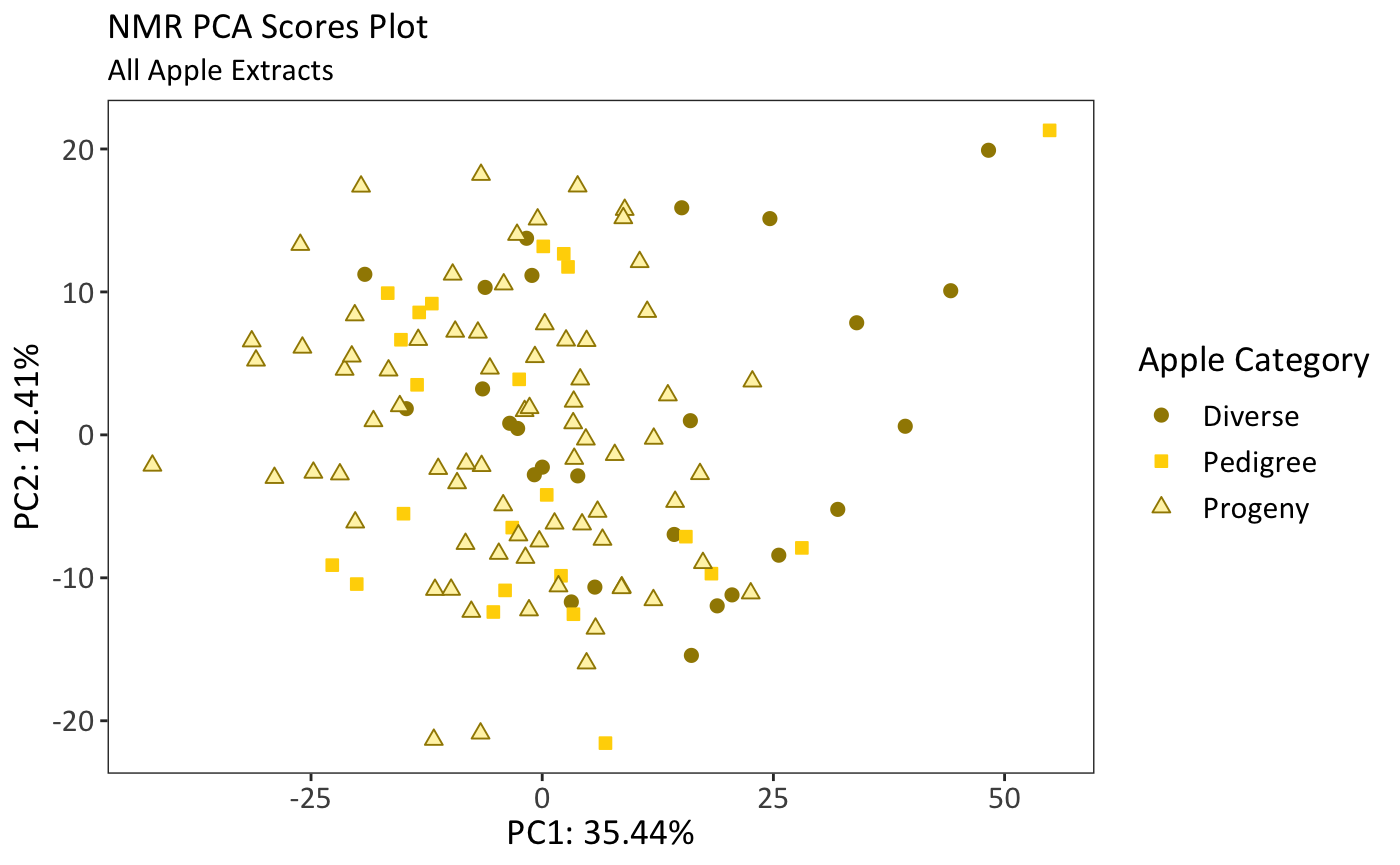

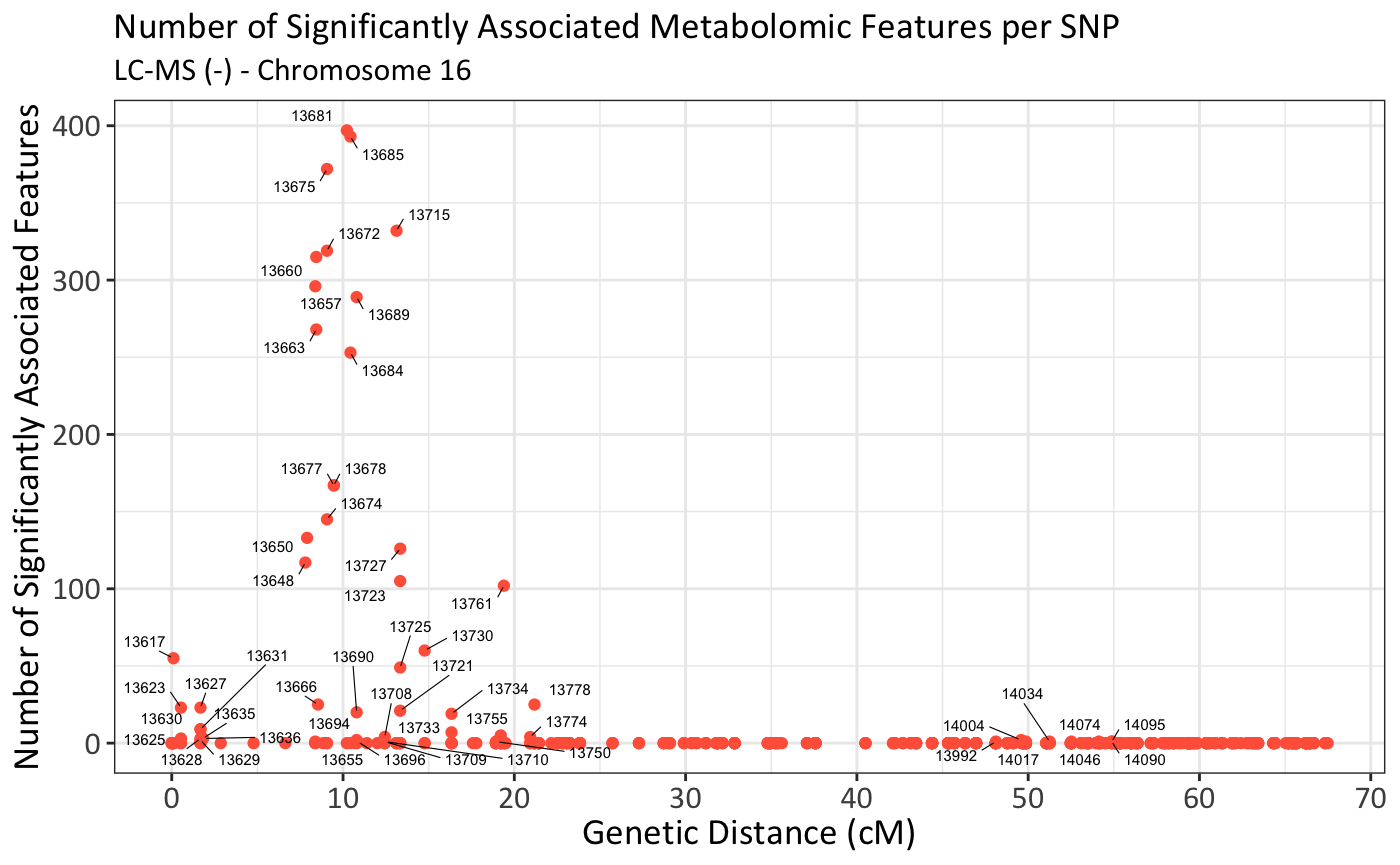

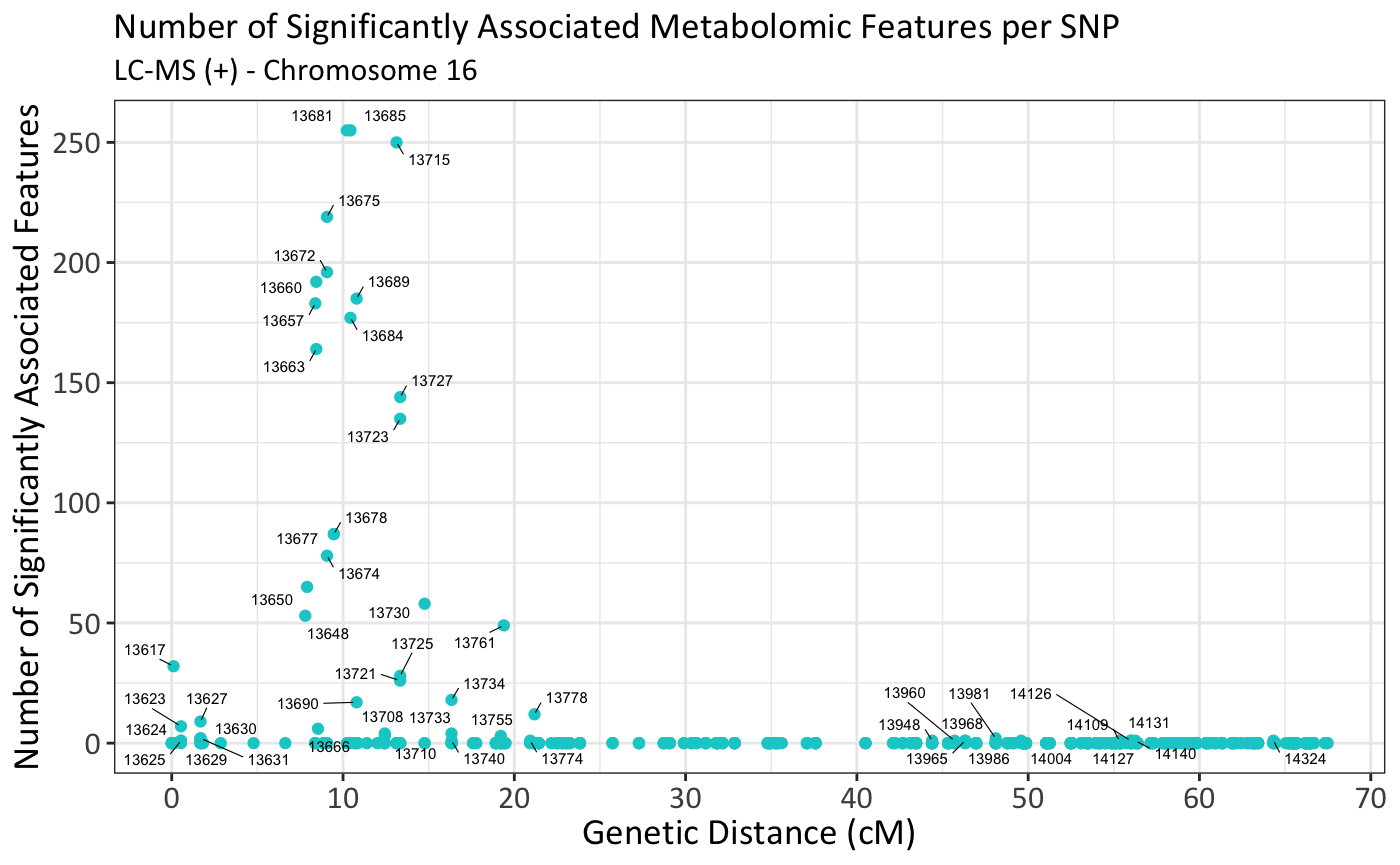
**Fig. S6** Plots displaying the number of significantly associated metabolomic features per SNP on chromosome 16. Significance is determined as -log(*p*) ≥ 5 for LC-MS data sets and ≥ 4 for NMR. SNPs are plotted based on their genetic position (cM) on the x-axis. SNPs with a minimum of 1 significant feature association are labeled with their study index number. Top SNPs include: 13681(SNP_FB_1074682), 13685 (RosBREEDSNP_SNP_CT_1540624_Lg16_LAR1_MAF40_1618769_exon2), and 13675 (SNP_FB_0335535). Additional index-to-SNP name conversions, including synonyms from the Affymetrix 480K SNP array, are available in Table **S10**.

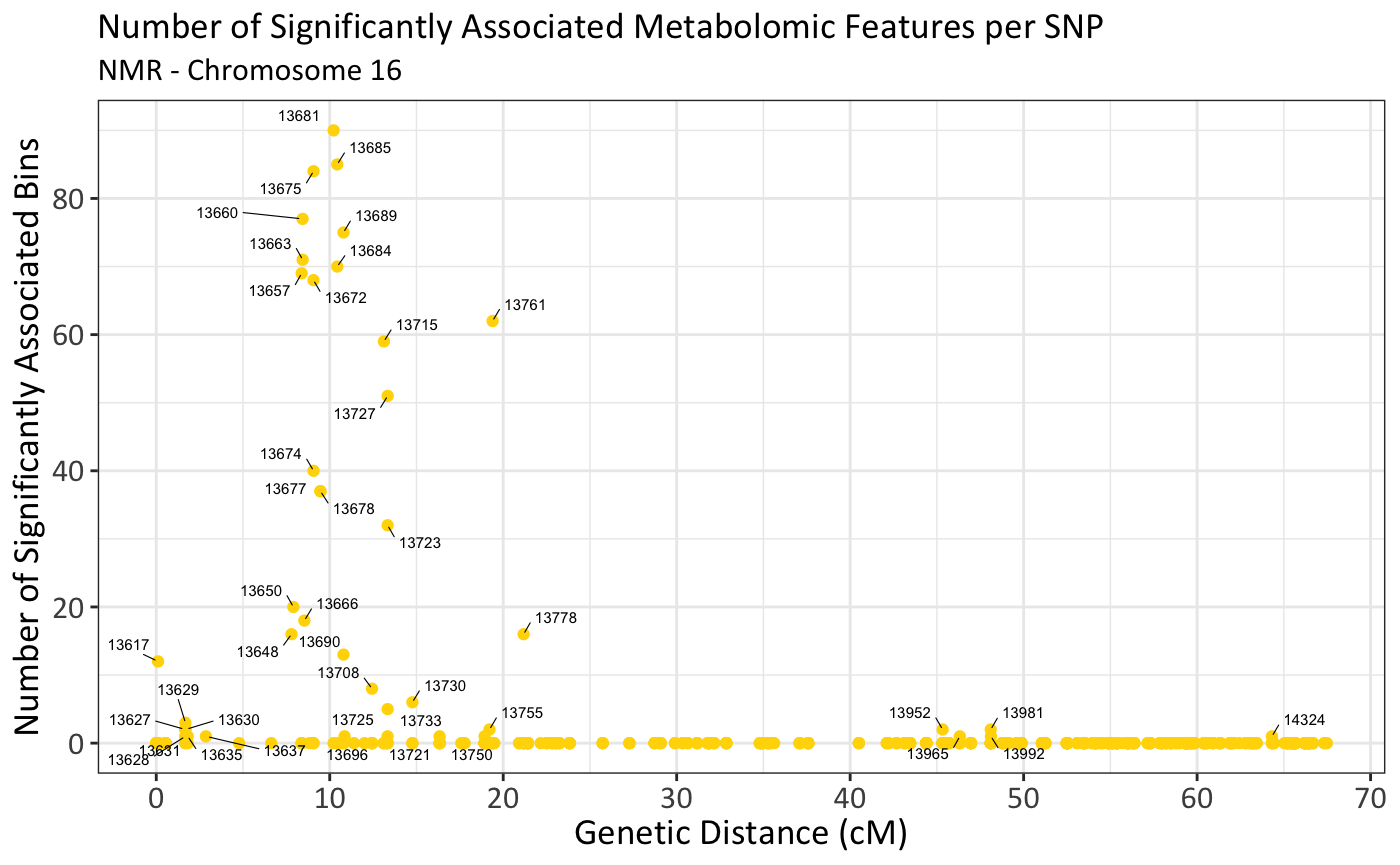

**Fig. S7** Plots displaying the number of significantly associated metabolomic features per SNP on chromosome 17. Significance is determined as -log(*p*) ≥ 5 for LC-MS data sets and ≥ 4 for NMR. SNPs are plotted based on their genetic position (cM) on the x-axis. SNPs with a minimum of 1 significant feature association are labeled with their study index number. Top SNPs include: 15109 (RosBREEDSNP_SNP_AG_20028330_Lg17_01298_MAF50_1664885_exon1), 15123 (SNP_FB_1114677), and 15133 (SNP_FB_0398770). Additional index-to-SNP name conversions, including synonyms from the Affymetrix 480K SNP array, are available in Table **S10**.

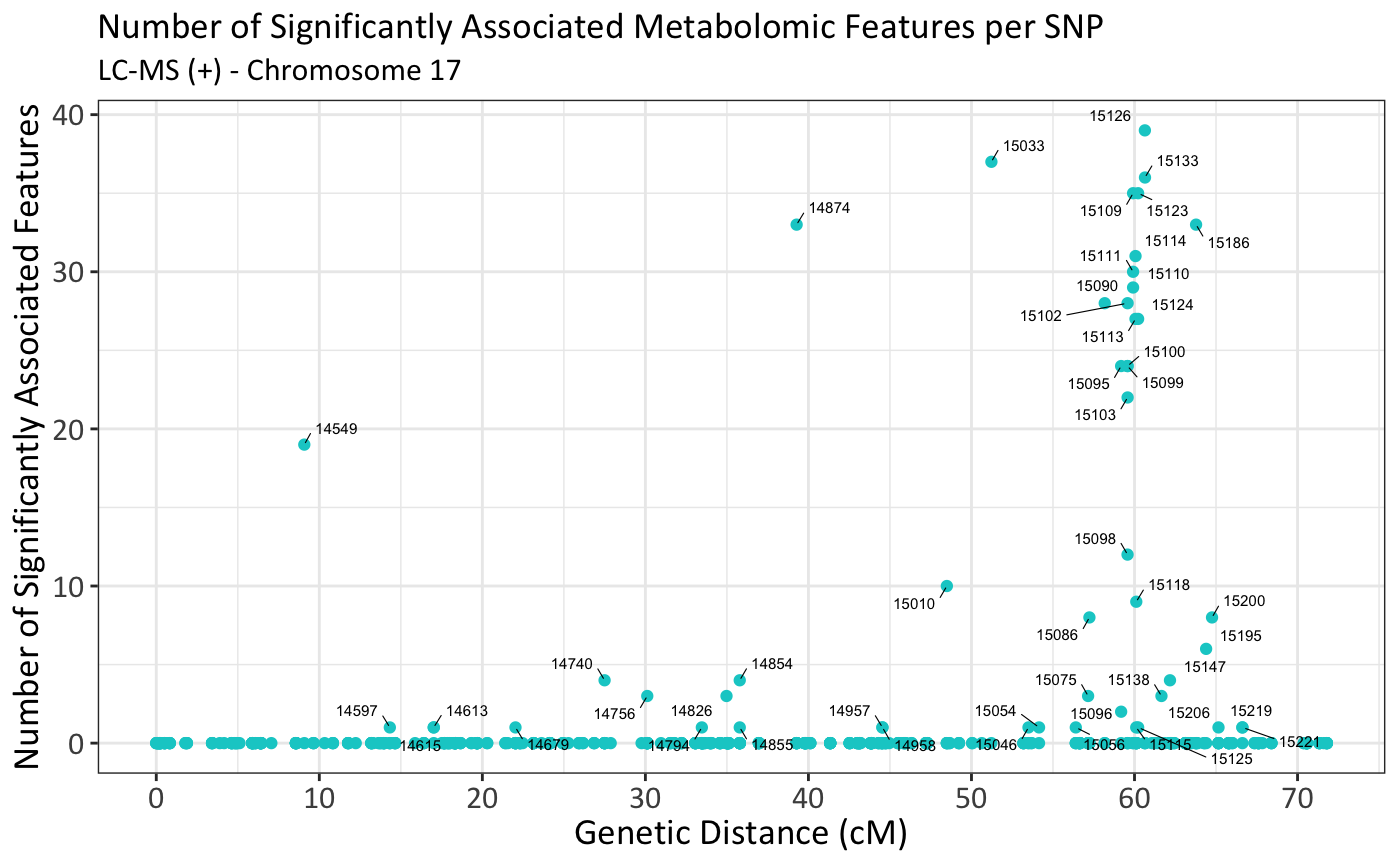

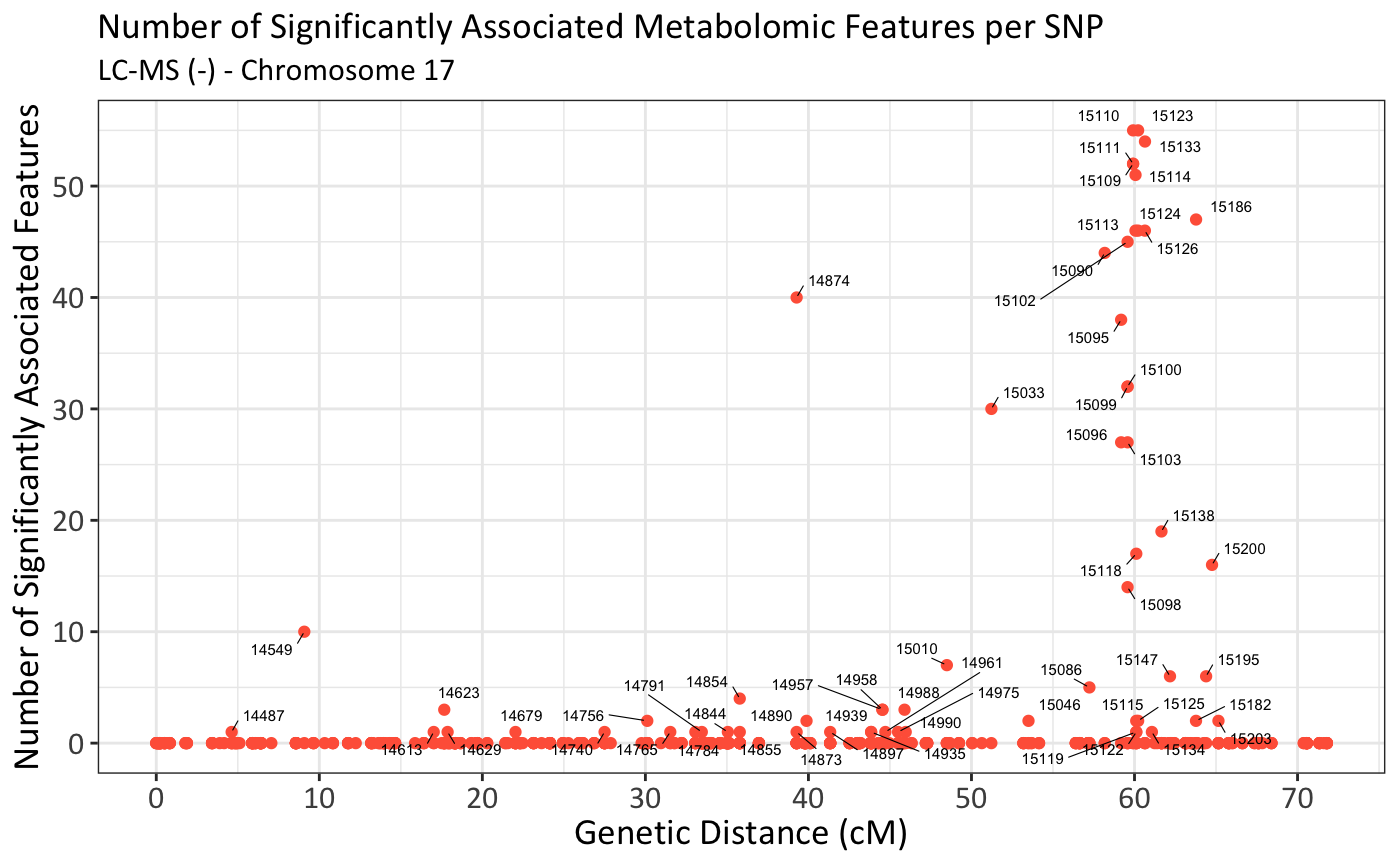

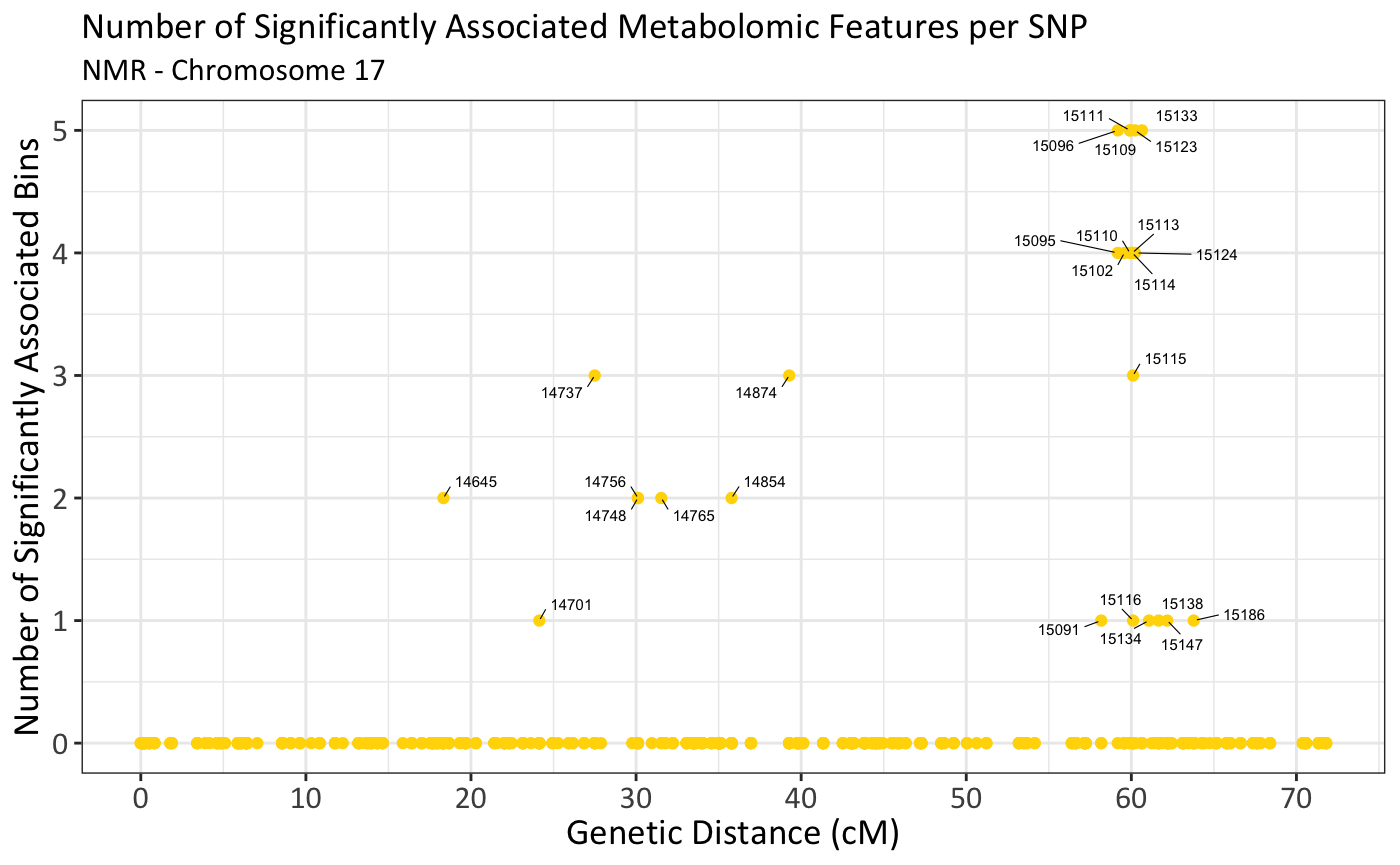

**Fig. S8** 1D ^1^H NMR spectrum of the apple extract pooled quality control. Yellow lines indicate each bin that was significantly associated with at least one SNP. Dashed lines approximately divide the spectrum according to the type of compounds that elicit peaks at that chemical shift. The aromatic region and amino acid region are in much lower abundance than the sugar region, so magnified inserts are also presented.

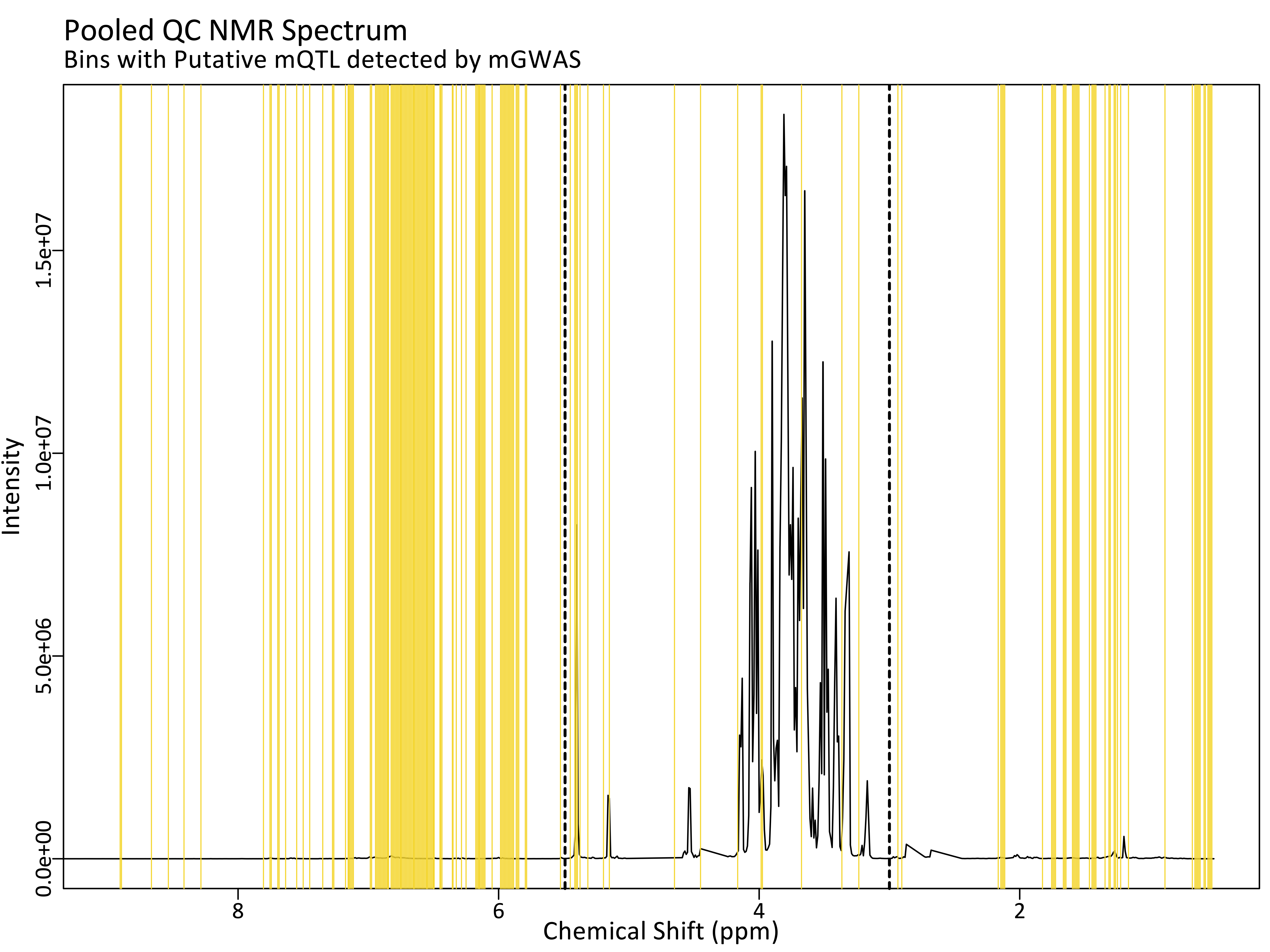

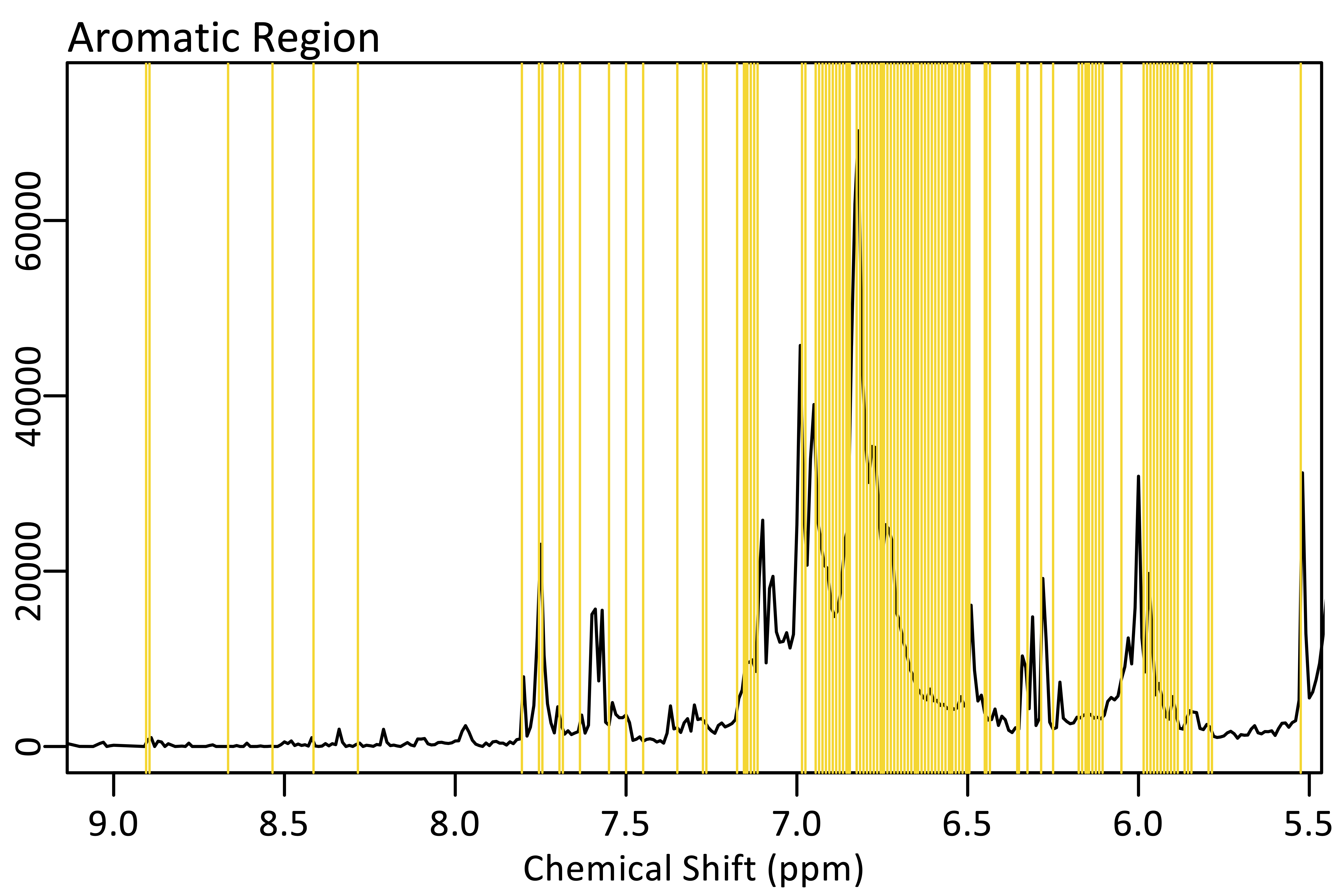

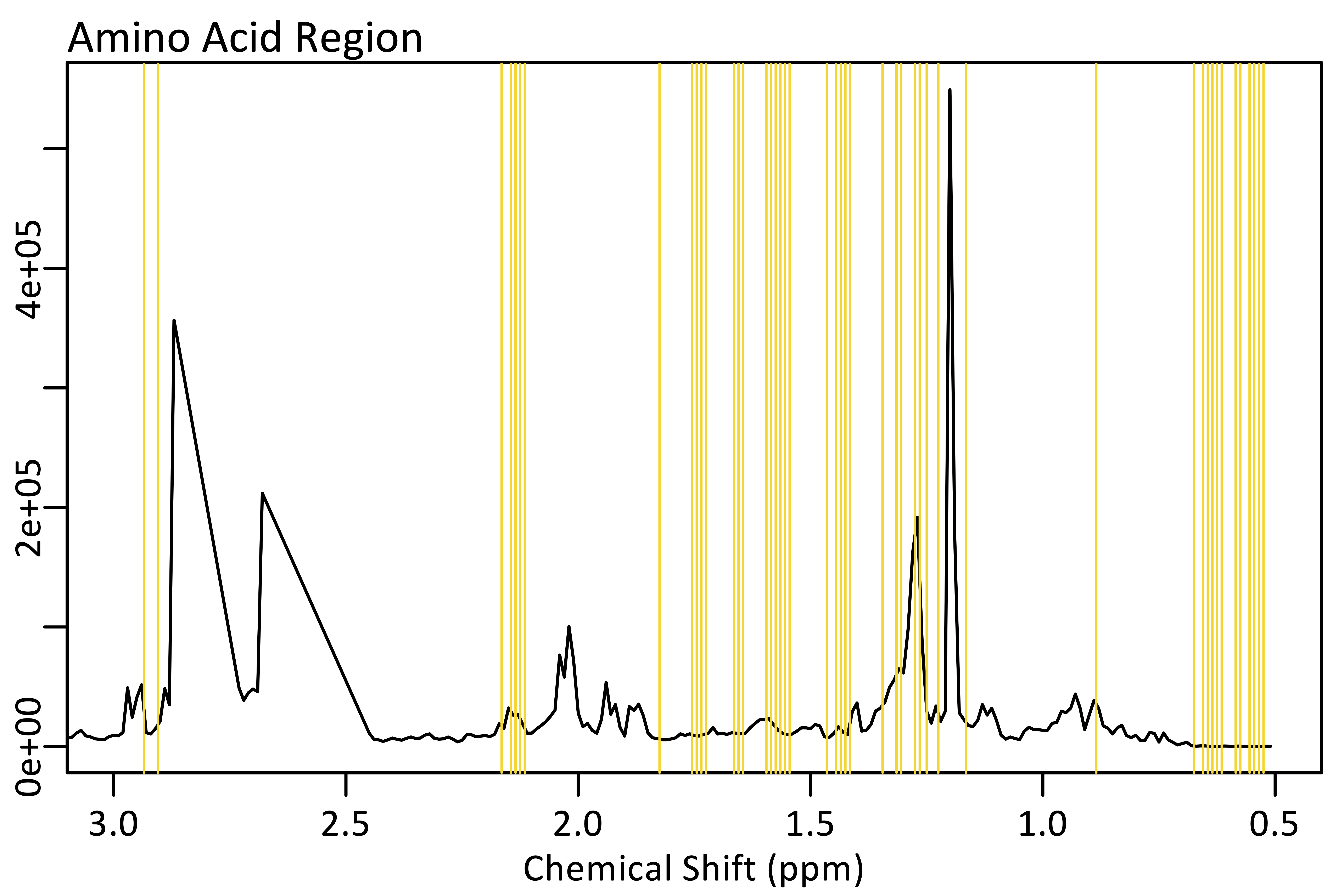

Sugar Region

**Table S1** Complete metadata compilation for all samples in an Excel spreadsheet

**Table S2** SNP data used in mGWAS

**Table S3** Instrument protocol for dry down of apple extracts for NMR analysis.

**Table S4** Parameters set for full scan LC-MS positive ionization mode data deconvolution in mzMine2.51.

**Table S5** Parameters set for full scan LC-MS negative ionization mode data deconvolution in mzMine2.51.

**Table S6** Deconvoluted data matrix of features and samples from full scan LC-MS positive ionization mode analysis.

**Table S7** Deconvoluted data matrix of features and samples from full scan LC-MS negative ionization mode analysis.

**Table S8** Processed data matrix of bins (0.01 ppm) and samples from 1D 1H NMR spectroscopy analysis.

**Table S10** SNP index numbers for our study along with the SNP name for the 20K SNP array, synonyms for the 480K Affymetrix SNP array, linkage group, genetic distance (cM) according to the iGLMap.

**Table S11** Parentage information used in calculating the A matrix using R package AGHmatrix. Columns are Individual, Parent 1, and Parent 2. “0” indicates unknown/uninformative parentage.

**Table S12** SNP data used for calculating the G matrix using R package AGHmatrix. Rows are apple genotypes/individuals, and columns are SNPs. Pedigree-related individuals are included for which there was no metabolomics data collected, but SNP data was available. The 0, 1, 2 scheme is used as required by the package. “NA” indicates missing data.

**Table S13** Output from mGWAS in the form of -log10(P) values for each SNP-metabolomic feature combination for the analysis of LC-MS (+) data of the Diverse population set.

**Table S14** Output from mGWAS in the form of -log10(P) values for each SNP-metabolomic feature combination for the analysis of LC-MS (+) data of the Pedigree population set.

**Table S15** Output from mGWAS in the form of -log10(P) values for each SNP-metabolomic feature combination for the analysis of LC-MS (+) data of the Progeny population set.

**Table S16** Output from mGWAS in the form of -log10(P) values for each SNP-metabolomic feature combination for the analysis of LC-MS (-) data of the Diverse population set.

**Table S17** Output from mGWAS in the form of -log10(P) values for each SNP-metabolomic feature combination for the analysis of LC-MS (-) data of the Pedigree population set.

**Table S18** Output from mGWAS in the form of -log10(P) values for each SNP-metabolomic feature combination for the analysis of LC-MS (-) data of the Progeny population set.

**Table S19** Output from mGWAS in the form of -log10(P) values for each SNP-metabolomic feature combination for the analysis of NMR data of the Diverse population set.

**Table S20** Output from mGWAS in the form of -log10(P) values for each SNP-metabolomic feature combination for the analysis of NMR data of the Pedigree population set.

**Table S21** Output from mGWAS in the form of -log10(P) values for each SNP-metabolomic feature combination for the analysis of NMR data of the Progeny population set.

**Table S22** A list of metabolomic features for LC-MS (+), (-), and NMR that remained in the pipeline after filtering for significance and intersecting for significance across the three population sets. LC-MS features are given as m/z and retention time. NMR bins are given as the 0.01 ppm-wide chemical shift.

**Table S23** Summarized output from the targeted pedigree-based analysis in FlexQTL™ for the breeding parents and progenies only. The data are organized by family to visualize haplotypes within the progenies. The haplotypes are coupled with the log2 chlorogenic acid abundance from LC-MS (+), breeding value, assigned mQTL genotype (i.e., QQ, Qq, qq), and family designation.

**Table S24** Summarized output from the targeted pedigree-based analysis in FlexQTL™ for all pedigree-related individuals. The haplotypes are coupled with the log2 chlorogenic acid abundance from LC-MS (+), breeding value, and assigned mQTL genotype (i.e., QQ, Qq, qq).

**Table S25** Chlorogenic acid mQTL Candidate genes of the phenylpropanoid pathway and their associated metadata including enzyme name, acronym, EC number, gene name for Malus domestica, and physical location on chromosome 17.

**Methods S1** Apple fruit and leaf collection details

To ensure the breeding-relevance of our germplasm, the majority of apples for the study (n=147) were collected from one of three commercial orchards with membership in the Midwest Apple Improvement Association (MAIA) grower-participatory breeding program: Lynd Fruit Farm, Johnstown, OH (n=28); Whitehouse Fruit Farm, Canfield, OH (n=68); and David Doud’s Countyline Orchard, Wabash, IN (n=39) (Table **S1**). Additional varieties were obtained from Purdue University in West Lafayette, IN, (n=4); the USDA Apple Germplasm Repository in Geneva, NY, (n=2); The Dawes Arboretum in Newark, OH, (n=5); or other locations in Ohio (n=1) (Table **S1**).

Leaf tissue and apple fruit were collected from one tree for the majority of varieties included in the study with a few exceptions. Four apple varieties (‘Honeycrisp’, ‘Goldrush’, ‘Golden Delicious’, and ‘EverCrisp’) were collected at each of the three MAIA sites. Also, 13 of the 27 ‘Honeycrisp’ × ’Fuji’ progeny from David Doud’s Countyline Orchard in Wabash, IN, were also collected from clonal replicates at White House Fruit Farm in Canfield, OH. Phenotypic data for varieties with fruit collection at multiple sites was averaged for subsequent analysis. Leaf tissue was collected in summer 2018 for DNA extraction. Fruits were harvested in fall 2018 for metabolomic analysis (Table **S1**).

A minimum of three apples were collected from each selected tree when optimally ripe, as determined by expert opinion based on ground color and traditional ripening timeline for specific varieties. All samples were sent to the Ohio Agricultural Research and Development Center in Wooster, OH. Apples were stored in a cooler at 4 °C for no more than seven days. Three apples were rinsed, dried, and cored for each selection. Eight slices were chosen at random then flash frozen in liquid nitrogen. To ensure adequate sample quantity, more than three fruits were cored and frozen if fruits were small. Fruits too small to be cored were cut with a knife or frozen whole. The frozen slices were placed in labeled freezer storage bags and stored at -20 °C until extraction. Comprehensive metadata concerning the samples are available in Table **S1**.

**Methods S2** Methanolic apple fruit extraction process and metabolomics sample preparation

Solvents and chemicals were purchased from Fisher Scientific (Pittsburg, PA, USA), and metabolomics solvents were LC-MS/Optima grade. Authentic standards were purchased from Sigma-Aldrich (St. Louis, MO, USA).

A total of 147 unique apple samples were prepared in random order for metabolomic analysis. Several representative slices were removed from each sample freezer bag. For each apple sample, peel was removed and weighed to 1.0 g ± 0.05 g. Pieces of apple flesh were cut from the slices and added to the peel to reach a combined weight of 5.0 g ± 0.05 g. One sample (*M. floribunda*) with inadequate sample collection was weighed to half the value of the standard: peel 0.5 g ± 0.05 g and flesh 2.5 g ± 0.05 g. Separate weighing of peel and flesh was used to account for the different sizes of the apples. Small apples would have a higher peel:flesh ratio than larger apples. This would be problematic in extraction because there is a higher concentration of phytochemicals in the apple peel compared to flesh. Weight matching peel and flesh separately allowed us to see differences in chemical abundance based on true differences instead of differences imparted by fruit size.

Weight-matched peel and flesh were then placed together into tube with two 3/8″ × 7/8″ angled ceramic cutting stones (W.W. Grainger: Lake Forest, IL, USA; Item no.: 5UJX2) and 15 mL of methanol to extract polar/semi-polar metabolites and inhibit enzymatic and non-enzymatic oxidation reactions. After overnight storage at -80°C, tubes were placed in a sample homogenizer (SPEX® SamplePrep Geno/Grinder®, NJ, USA) for grinding. Samples were then centrifuged at 2,800 × *g* for 3 minutes to pellet insoluble material. Supernatant was then syringe-filtered (0.22 μm PTFE) to remove remaining particulates. Filtered extract was then dispensed for UHPLC-QTOF-MS and NMR analyses. Samples for LC-MS analysis were diluted to 50:50 water:methanol. NMR samples were dried via a vortex vacuum evaporator (Combidancer, Hettich AG, Baech Switzerland) (Table **S3**).

A 1.0 mL aliquot of each sample extract was pooled to create a bulk quality control (QC) to be used in the MS experiments. The pooled QC solution was diluted with H_2_O to 50% MeOH then aliquoted. Several process blanks were made by conducting the extraction method with all the same physical elements except the apple fruit. Analyzing the process blank allowed detection of the presence of any residues or contamination with compounds from the pipette tips, tubes, or pellets used in the extractions. All extracts were stored at -20 °C until analysis.

At the time of NMR analysis, dried samples were reconstituted using an 80:20 solvent system of deuterated methanol (CD_3_OD) and 0.2 mM aqueous phosphate buffer. Trimethylsilylpropanoic acid (TSP) was added to the bulk solvent to serve as an internal reference for chemical shift. For each dried sample, 800 μL of the solvent system was added. Samples were then vortexed to help dissolve extract and placed in a sonication water bath at room temperature for 5 minutes. For each sample, 600 μL was then transferred to a 5 mm NMR tube.

**Methods S3** Full scan UHPLC-ESI-QTOF-MS method details, instrument settings, and data deconvolution

Polar extracts were analyzed with an Agilent 1290 Infinity II series UHPLC coupled to an Agilent 6545 quadrupole time of flight mass spectrometer with electrospray ionization (ESI-QTOF-MS) (Agilent, Santa Clara, CA, USA). Injection volume was 3 μL per sample. Reverse phase chromatography was conducted with a Waters Acuity UPLC HSS T3 column (2.1 x 50 mm, 1.8 μm particle size) maintained at 40 **°**C (Waters, Milford, MA, USA). Full-scan spectral data was collected separately in both positive and negative ionization modes for complete metabolite coverage. Mobile phases consisted of water with 0.1% formic acid (A) and acetonitrile with 0.1% formic acid (B). Flowing constantly at 0.5 mL/min, the gradient was as follows: 0-0.5 min 0% B; 0.5-8 min increase to 100% B; 8-9 min hold 100% B; 9.01-10.0 isocratic at 0% B for a total run time of 10 minutes. MS parameters were as follows: gas temp 350 **°**C, gas flow 10 L/min, nebulizer 35 psig, sheath gas temp 375 **°**C, sheath gas flow 11 L/min, VCap 4500 V, nozzle voltage 500 V, fragmentor 100, skimmer1 45, octopoleRFPeak 750, and scan rate 2 spectra/s with a mass range of 100-1700 *m/z*.

Before injecting any apple extracts, solvent blanks (1:1 water:methanol) were injected to equilibrate the instrument. Next, process blanks were run to give a baseline of chemical noise present in the instrument and from the extraction process. Pooled QCs were then injected repeatedly until base peak chromatograms were consistent. Then, order of sample analysis followed the random order established by sample weighing so that samples which were extracted first were also analyzed first. Identical pooled QCs were analyzed every seventh injection, providing a way to monitor instrument stability and data quality across the experiments.

Raw spectral data were converted to *.mzML files using msConvert in ProteoWizard with 64-bit encoding precision without zlib compression (Chambers, MacLean, Burke, *et al.*, 2012). Data was then deconvoluted in MZmine2.51 (Pluskal, Castillo, Villar-Briones, *et al.*, 2010) with a pipeline including mass detection using wavelets (ADAP), Automated Data Analysis Pipeline (ADAP) chromatogram building (Myers, Sumner, Li, *et al.*, 2017), feature detection, isotope grouping, alignment, gap filling, and filtering according to various parameters recorded in Table **S4-S5**. This produced a data matrix of signal intensities, retention time, and *m*/*z* ratio for each feature in each sample. Manual data cleanup included removal of features with intensity less than 10 times that detected in the process blank and those with >30% coefficient of variance across the pooled QC samples. Intensities were doubled for the sample that was weighed to half the target weight for peel and flesh. For each remaining feature, missing values were imputed with half the minimum intensity.

**Methods S4** 1D 1H NMR spectroscopy method, instrument settings, and data processing

1D ^1^H NMR experiments were performed using a Brüker Avance III spectrometer (Bruker, Ettlingen, DE), operating at 700.13 MHz, equipped with a TXO helium-cooled 5 mm triple-resonance observe probe with samples at 25 ± 0.1 °C. Spectra were acquired using the noesypr1d pulse sequence with the following parameters: 64 scans and 4 dummy scans, 64K data points, 90° pulse angle (10.5 μs), relaxation delay 3 s, spectral width 15 ppm. The spectra were acquired without spinning the NMR tube in order to avoid artifacts, such as spinning side bands of the first or higher order.

The spectra were processed using the Topspin software package provided by Brüker Biospin. A polynomial fourth-order function was applied for baseline correction in order to achieve accurate quantitative measurements when integrating signals of interest. Phase correction and baseline correction were adjusted manually. Chemical shifts are reported in ppm from TSP (δ=0).

Post-processing of spectra was performed using R package *mrbin* v1.2.9001 (Klein, 2020). Spectra were binned with a bucket size of 0.01 ppm across the range of the spectrum (9.5-0.5 ppm). Regions for water (5.0-4.6 ppm) and methanol (3.335-3.305 ppm) peaks were excluded. Three unstable peaks affected by slight pH shifts were each summed (2.68-2.45, 2.87-2.73, 4.45-4.25). Negative intensities were converted to positive values with an affine transformation. Bins were filtered with a noise threshold of 0.01 and a signal-to-noise-ratio (SNR) of 10. Intensities were doubled for the sample that were weighed to half the target weight for peel and flesh.

**Methods S5** Data-dependent UHPLC-ESI-QTOF-MS/MS method details, instrument settings, and data processing

Using the same gradient and QTOF settings as above, the pooled QC samples were also analyzed with MS/MS experiments (MassHunter B09 acquisition software, Agilent Technologies) in which data-dependent MS/MS spectra were collected using Agilent’s “iterative” MS/MS process. The goal was to collect MS/MS spectra of as many features from the apple extracts as possible for use in manual compound identification and molecular networking using Global Natural Products Social Molecular Networking (GNPS) (Wang, Carver, Phelan, *et al.*, 2016), both based on fragmentation. To balance between collecting both comprehensive and good quality spectra, the same QC sample was injected repeatedly, and the most abundant ions throughout the chromatogram were triggered for MS/MS data collection. Those ions were automatically added to an exclusion list, such that the next most abundant ions can then be selected for MS/MS. This allowed collection of fragmentation data for features across a range of ion intensities. These experiments were conducted in both positive and negative ionization mode with five injections for each of two collision energies, 20 and 40 eV.

MS settings were as above in full scan experiments. Parameters for AutoMS2 scans were as follows: MS minimum range 40 *m/z*, MS maximum range 1700 *m/z*, MS scan rate 3 spectra/s, MS/MS minimum range 40 *m/z*, MS/MS maximum range 1700 *m/z*, MS/MS scan rate 1 spectra/s, isolation width narrow (~1.3 amu), and decision engine advanced. Terms for precursor selection were max precursors per cycle 2, threshold (absolute) 10,000, threshold (relative)(%) 0.100, precursor abundance based scan speed – yes, target 100,000 counts/spectrum, use MS/MS accumulation time limit – no, use dynamic precursor rejection – no, purity stringency 100%, purity cutoff 30%, common isotope model, active exclusion enabled – yes, active exclusion excluded after 2 spectra, active exclusion released after 0.12 min, and sort precursors by abundance only.

**Methods S6** Feature identification based on online databases and data-dependent MS/MS spectral matches and classical molecular networking in GNPS.

Databases, such as the Human Metabolome Database (HMDB) (Wishart, Feunang, Marcu, *et al.*, 2018), were searched for potential matches for *m/z* values for features of interest. If fragmentation data had been obtained in the untargeted data-dependent MS/MS experiments, MS/MS spectral data were also compared against experimental or predicted MS/MS spectra in databases as well as fragmentation data or spectra in the literature.

Generation of putative identities was broadly applied to all collected iterative MS/MS spectra using the classical molecular networking platform in the GNPS infrastructure (Wang, Carver, Phelan, *et al.*, 2016). A molecular network was created using the online workflow (https://ccms-ucsd.github.io/GNPSDocumentation/) on the GNPS website (http://gnps.ucsd.edu). The precursor ion mass tolerance was set to .01 Da and a MS/MS fragment ion tolerance of 0.01 Da. A network was then created where edges were filtered to have a cosine score above 0.65 and more than 6 matched peaks. Further, edges between two nodes were kept in the network if and only if each of the nodes appeared in each other's respective top 10 most similar nodes. Finally, the maximum size of a molecular family was set to 100, and the lowest scoring edges were removed from molecular families until the molecular family size was below this threshold. The spectra in the network were then searched against GNPS' spectral libraries. All matches kept between network spectra and library spectra were required to have a score above 0.7 and at least 6 matched peaks. This workflow was repeated as four separate jobs—one per ionization mode, collision energy combination: negative 20 eV, negative 40 eV, positive 20 eV, and positive 40 eV.

Chlorogenic acid stock solution was prepared in 1:1 MeOH:H_2_O. Stock solutions were analyzed with full scan LC-MS and MS/MS with collision energies of 20 and 40 eV. Fragments of note were 191.0574 *m/z* and 161.0247 *m/z*, which corresponded to the constituent molecules of chlorogenic acid, 3-*O*-caffeoylquinic acid: quinic acid ([M-H]^-^ 191.0562 *m/z*) and caffeic acid ([M-H_2_O-H]^-^ 161.0239 *m/z*), respectively. Identities of NMR peaks from the complex mixture were investigated primarily through comparison with identities determined in the literature for polar apple extracts as well as other fruits. Chemical shift and multiplicity were also compared to online databases containing 1D ^1^H NMR spectral data for pure compounds, namely HMDB (Wishart, Feunang, Marcu, *et al.*, 2018). For compound ID verification in NMR for chlorogenic acid, spike experiments with the authentic standard were analyzed with additional 1D ^1^H NMR analysis. Here, the analytical standard was added to a pooled QC sample and analyzed to determine if the intensity increases for the peaks of interest at specific ppm.

**Methods S7** Details concerning pedigree-based analyses (PBA) under a Bayesian framework conducted in FlexQTL™.

The effect of environment was obviated from the analyses, as it was not practical to increase the model complexity. A minimum of 50,000 Markov chain Monte Carlo sweeps were run for each routine, thinning every 50 iterations with a thin screen of 1,000. The minimum effective sample size (ESS) to consider that convergence was achieved in the model was set to 101. If at 50,000 iterations, ESS < 101, the software continued iterations, up to 100,000, to ensure convergence and sound statistical foundation for findings. The Finite Polygenic Model was also run to identify major (Mendelian) genes. The algorithms were applied to the data within at the OSC (OSC 1987). Three replications were conducted for each trait input to provide additional certainty in estimations and statistical robustness for later replication of the results by others.
